# Introgression of a block of genome under infinitesimal selection

**DOI:** 10.1101/227082

**Authors:** Himani Sachdeva, Nicholas H. Barton

## Abstract

Adaptive introgression is common in nature and can be driven by selection acting on multiple, linked genes. We explore the effects of polygenic selection on introgression under the infinitesimal model with linkage. This model assumes that the introgressing block has an effectively infinite number of genes, each with an infinitesimal effect on the trait under selection. The block is assumed to introgress under directional selection within a native population that is genetically homogeneous. We use individual-based simulations and a branching process approximation to compute various statistics of the introgressing block, and explore how these depend on parameters such as the map length and initial trait value associated with the introgressing block, the genetic variability along the block, and the strength of selection. Our results show that the introgression dynamics of a block under infinitesimal selection is qualitatively different from the dynamics of neutral introgression. We also find that in the long run, surviving descendant blocks are likely to have intermediate lengths, and clarify how the length is shaped by the interplay between linkage and infinitesimal selection. Our results suggest that it may be difficult to distinguish introgression of single loci from that of genomic blocks with multiple, tightly linked and weakly selected loci.

---

Limited intogression of genetic material between closely related sub-species or species is common, even among species displaying significant divergence (Arnold 2004; Hedrick 2013; Racimo et al 2015). Introgression may be adaptive if it supplies new genetic variation that facilitates a response to selection. Well-documented examples include adaptive introgression between different species of sunflowers, resulting in increased herbivore resistance in the recipient species (Whitney et al. 2006), introgression between the Algerian mouse and the European house mouse, which most probably caused the latter to acquire increased pesticide resistance (Song et al. 2011), and the possible introgression between Denisovans and the ancestors of modern-day Tibetans, resulting in altitude adaptations in Tibetan populations (Huerta-Sanchez et al. 2014).

Unlike de novo mutation, migration introduces multiple, linked allelic variants into a population. The likelihood of successful introgression or hybridization is thus sensitive to the density of selected loci among these variants and the distribution of their fitness effects, in addition to demographic parameters (Yeaman and Whitlock 2011). More generally, it has been argued that *polygenic* adaptation or introgression is likely to involve minor shifts in allele frequencies as opposed to selective sweeps, leading to genomic signatures that are qualitatively distinct from those of major-effect loci (Pritchard et al 2010). Similarly, background selection due to many weakly deleterious loci is expected to shape diversity at linked sites differently from a single strongly deleterious locus (Good et al. 2014). Incorporating linked, polygenic selection into methods detecting introgression is thus an important challenge for population genetic inference (Elyashiv et al 2016).

A particularly useful limit for studying the dynamics of multiple selected loci is the infinitesimal model which assumes that a given phenotype is influenced by a very large (effectively infinite) number of unlinked or weakly linked loci with infinitesimally small effects (Bulmer 1980; Barton et al. 2017). It follows that the effect of selection on the allele frequency at any one locus is negligible, so that the genic variance can be assumed to be constant over short time scales (in the absence of drift). The response to selection is then primarily via an increase in the genetic variance of the population, which is driven by changes in the linkage disequilibria (LD) between loci (Bulmer 1980).

The assumption of unlinked loci is, however, problematic if the density of selected variants on the genome is very high. To explore the effect of linked selection on introgression, we consider a trait determined by an effectively infinite number of loci but assume that these are now uniformly distributed on a genomic block of map length *y*_0_. This model, which we refer to as the ‘infinitesimal model with linkage’ was first introduced by Robertson (1977). It is parameterized by a single parameter *V*_0_, the genic variance per unit map length (discussed in more detail below).

For simplicity, we consider a scenario where a single block of genome from a source population enters a *genetically homogeneous* native population having zero genic variance. By definition, the native population also has zero *segregation variance*, which is the variance in trait values of offspring produced by mating between any two randomly chosen individuals in the population. This implies that any adaptive response must be based purely on the genetic variation supplied by the ‘introduced genome’. We typically follow descendants of the introduced genome over a few hundred generations after the initial hybridization event and neglect the creation of new variation in the native population by mutation over these time scales.

The assumption of a genetically homogeneous native population is unrealistic, but may approximately describe a situation where the native population has a small effective population size and is subject to strong stabilizing selection which depletes polymorphisms, resulting in much lower segregation variance than in the source population. A shift in the selection optimum at a later time would then provide further opportunities for evolution when new variation is introduced via migration. The main advantage of considering a genetically homogeneous native population is that it allows us to focus purely on the dynamics of the introduced block, without considering additional effects that arise due to the association of this block (or its descendants) with different native genomic backgrounds.

The introgression of a neutral block of genome was modeled as a branching process by Baird et al. (2003). They found that a neutral block of map length *y* is lost very slowly from the population— the probability that at least some portion of the block survives at time *t* falls as ~*y*/ log(*yt*/2), in contrast to the faster ~1/*t* decay observed for a single neutral locus. This is explained by the fact that a large block has a higher chance of being transmitted to offspring than a small block (either as a whole or in part), resulting in a relatively large number of descendants carrying at least a part of the block, in the first few generations. Over longer time scales, as surviving fragments become smaller and smaller, this transmission advantage is lost. However, by this time there are so many (~*yt*) of these small descendant blocks, that the probability of all of them being lost from the population becomes very small.

Baird et al’s analysis does not include selection per se, though it can be used to describe a rather artificial situation in which any fragment of the block, howsoever small, is associated with the same selective effect. However, in the infinitesimal limit, we expect the additive effects of short fragments to be smaller than those associated with long fragments (on average). Thus, recombination not only splits the original block into smaller and smaller fragments, but also dilutes the selective (dis-)advantage of descendants carrying fragments of the block. This can result in a complex feedback whereby the strength of recombination relative to selection shapes the distribution of block sizes and trait values, while this distribution itself determines the effective strengths of selection and recombination in the population.

The main goal of our work is to use the infinitesimal framework to understand how the resultant, *dynamically changing* balance between selection and recombination influences the introgression of genomic blocks. An important focus is to clarify how the introgression of an extended block containing a very large number of loci with infinitesimally small selective effects differs from that of a single strongly selected locus. This also impinges on the broader question of whether weakly selected loci can have discernible effects on genome evolution under various selection-migration scenarios.

## 1 Model and Methods

We consider a scenario where a single haplotype from a diverged source population enters a diploid native population. Each individual in the native and source populations expresses a trait determined additively by a large number of small-effect genes, which, to a first approximation, are uniformly distributed over a genomic block of map length *y*_0_.

The native population is assumed to be genetically homogeneous and consist of individuals having *identical* haplotypes and hence the same trait value (*z*=0). For ease of analysis, we define the additive contributions of any allele (or alternatively, any small region) of the native haplotype to also be zero. The native haplotype thus provides a reference with respect to which the additive contributions of different tracts of the introduced block are measured.

The introduced block has an associated trait value *z*_0_, which is obtained by integrating over the contributions from different regions of the block (over a map length *y*_0_). These regions typically have unequal contributions to the trait value. Thus, when the introduced block is split by recombination, descendants inherit fragments of the block associated with a range of trait values, which differ from the trait value of the native haplotype (*z*=0) but also from each other.

The additive trait is under directional selection in the native population, such that individuals with trait value z have a Poisson-distributed number of offspring, with mean *w*(*z*)=2 exp(*βz*), where *β* represents the strength of selection. Thus, each individual carrying native haplotypes produces two offspring on average, while individuals carrying fragments of the introduced block may produce more or less than two, depending on the trait value associated with the fragment. This results in a response to selection, whose magnitude depends on the strength of selection *β*, the original trait value *z*_0_ associated with the introduced haplotype, as well as the extent of genetic variation among the descendants of this haplotype, which can be characterised quite simply within the infinitesimal framework described below.

### 1.1 Infinitesimal model with linkage and no selection

The infinitesimal model with linkage was introduced by Robertson (1977), and can be thought of as a limiting case of a trait determined additively by *D* discrete loci uniformly distributed on a genomic block of map length *y*_0_. Consider a population in linkage equilibrium (LE), in which the allelic states of different loci determining the trait are statistically independent of each other (irrespective of the extent of linkage between the loci). We further consider an individual block having trait value *z*_0_ within this population, and denote the additive contribution of the *i^th^* locus on this block by *α_i_*, such that 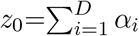. Then linkage equilibrium implies that *α_i_* are independent random variables drawn from some distribution (with variance *σ*^2^).

If we now adopt a coarse-grained view of this block, and consider the additive contributions of small genomic tracts of map length *δ*_*y*_0__ (instead of individual loci), then these are normally distributed with variance given by *~σ*^2^*D*(*δ*_*y*_0__/*y*_0_) for *δ*_*y*_0__≪*y*_0_. This follows simply from the central limit theorem under the assumption that the number of trait loci in a tract of length *δ*_*y*_0__, given by (*D*/*y*_0_)*δ*_*y*_0__, is sufficiently large, and that there is no LD in the population.

Thus the variance in additive contributions of small tracts of map length *δ*_*y*_0__ is proportional to *δ*_*y*_0__, with the constant of proportionality being *V*_0_=σ^2^(*D*/*y*_0_), which is the genic variance per locus (σ^2^) times the number of loci per unit map length (*D*/*y*_0_). We refer to *V*_0_ as the genic variance per unit map length, and note that the genic variance is just equal to the additive genetic variance minus the variance due to LD (which is zero here, under the assumption of LE).

It also follows that if we define *z*(*x*) as the trait value associated with the genomic region [0, *x*] of the block, then *z*(*x*) is a *Brownian path* (since there is no correlation or LD between additive contributions of different parts of the block). This Brownian path is characterised by two quantities— the trait value *z*_0_ associated with the full block (or the net displacement of the path in the interval [0,*y*_0_]), and the variance *V_0_* per unit map length (or the variance of the distribution from which increments to the Brownian path are drawn).

A key assumption of the infinitesimal model is that *V*_0_ is independent of the trait value *z*_0_, and is equal to the genic variance (per unit map length) in the source population. In Appendix S1 of the Supplementary Information (SI), we show that in the absence of selection, the variance of trait values in a diploid source population is simply 2*V*_0_*y*. Thus, in the infinitesimal framework, the parameter *V*_0_ measures the extent of variability along a single genome as well as the variability between genomes in the source population.

Based on the above model, we can write down the probability *R*(*y*_0_, *z*_0_→*y*_1_, *z*_1_) that given a block of map length *y*_0_ and trait value *z*_0_, the trait values associated with two constituent fragments of map length *y*_1_ and *y*_0_−*y*_1_ are *z*_1_ and *z*_0_−*z*_1_ respectively. This is just the joint probability distribution:

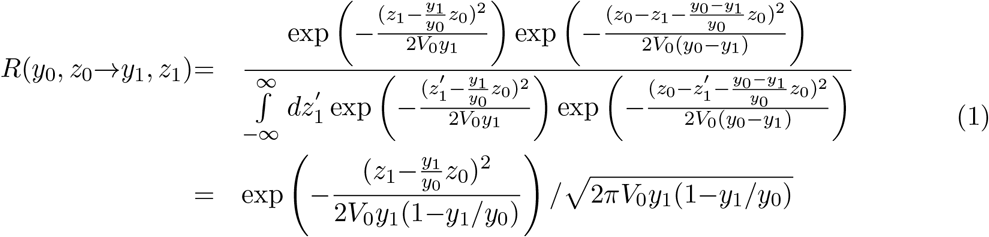

Thus, the trait value *z*_1_ associated with a daughter block (of map length *y*_1_) is normally distributed with mean (*y*_1_/*y*_0_)*z*_0_ and variance *V*_0_*y*_1_[1−(*y*_1_/*y*_0_)]. Equation (1) is the main equation describing inheritance under the infinitesimal model with linkage (see also p. 309, Robertson (1977)).

The infinitesimal model with linkage is more general than a commonly used selection model where the trait value (or fitness) of the descendant block is assumed to be *deterministically* proportional to its length, i.e., *z*_1_∝*y*_1_ (see for example, Barton (1983)). The deterministic selection model essentially describes a population that is fixed for equal effect alleles at all loci, and can be recovered by taking the *V*_0_→0 limit of the more general model described here.

Note that eq. (1) is not valid when the allelic states at physically linked loci are correlated. Such correlations may arise due to selection or genetic drift; stabilizing selection, in particular, builds up negative LD between genomic regions that are in tight linkage (Lande 1976). The variance released by recombination is then no longer described by eq. (1), but can be significantly lower than *V*_0_*y*_1_(1−*y*_1_/*y*_0_) for *y*_1_≪1. Thus any single genome encodes information about the nature of multi-locus associations in the source population from which it originates. This information lies hidden in the distribution of trait values of its constituent sub-blocks, and is revealed as recombination gradually isolates these sub-blocks.

We assume that there is no (or very weak) selection in the source population from which the ‘introduced haplotype’ originates. Then the source population is described by the infinitesimal model with linkage and no selection, while individual genomes within this population are represented by Brownian paths. In principle, we can generate this Brownian path during the course of one simulation run of the introgression process as follows— each time an individual passes on a fragment of the introduced block to a descendant, we decide on the trait value associated with the fragment by conditioning on the trait value of the parental block using eq. (1). We store trait values of different sub-blocks in a master list and keep updating this list with the values of smaller and smaller sub-blocks as they are are generated by recombination. While this scheme would exactly simulate the infinitesimal model with linkage, it is quite cumbersome to implement.

Instead, we follow an approximate simulation scheme which discretizes the Brownian path by dividing the introduced haplotype into *L*=2^*R*^ small sub-blocks of map length *δ*=*y*_0_/*L*. The additive contribution γ*i* associated with each sub-block is then chosen using an iterative scheme. First, the contributions of the two halves of the full block are set to *z* and *z*_0_−*z*, where *z* is sampled from the distribution given in eq. (1) with *y*_1_ set equal to *y*_0_/2. Then each of the two halves is further divided into two and the trait contributions of the resultant quarters chosen again using eq. (1). This process is iterated *R* times, thus obtaining additive contributions γ*i* of each of the *L*=2^*R*^ sub-blocks, ensuring that 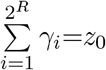. We try out various values of *R* and find that the initial dynamics of trait values is insensitive to the exact number of sub-divisions beyond *R*=10.

### 1.2 Introgression of a single block into an infinite, genetically homogeneous native population

To model the spread of the introduced haplotype within the native population, we make the following assumptions: (i) the map length *y*_0_ of the selected genomic region is small enough that multiple crossovers can be neglected (ii) descendants of the introduced haplotype form a small fraction of the population; thus, the likelihood of mating between two individuals, both bearing introgressed genetic material, is negligible. The second assumption is strictly valid only when the native population is infinite, but also holds approximately for a finite native population during the initial phase of introgression. Under the above assumptions, introgressed genetic material always appears in descendants as a single continuous block of map length *y*, surrounded by native blocks (with map length *y*_0_−*y*). Since the native population is assumed to be genetically homogeneous, it is sufficient to follow just the lengths and trait values of the fragments of the introduced block without considering the rest of the genome.

The first generation after the hybridization event is simulated by drawing the number of offspring of the individual carrying the introduced haplotype from a Poisson distribution with mean *w*(*z*_0_)=2exp(*βz*_0_), where *z*_0_ is the trait value associated with the introduced haplotype. Each of these offspring is either assigned no portion of the introduced block (with probability (1‒*y*_0_)/2), or the whole block (with probability (1−*y*_0_)/2), or a fragment of the block (with probability y_0_). The fragment is generated by choosing a single crossover point x that is uniformly distributed in the interval [0,*y*_0_], and then assigning to the offspring either the left [0,*x*] or the right [*x*, *y*_0_] fragment with equal probability. If the descendant block is smaller than the parent block, then the trait value of the descendant block is obtained by summing over the contributions of all the *complete* sub-blocks inherited from the parent plus a random contribution *ζ* representing the sub-block which is inherited in part. If the crossover point lies within the *i^th^* sub-block such that the descendant inherits a fraction *α* of this sub-block, then *ζ* is drawn from a normal distribution with mean equal to *α* times the additive contribution of the *i^th^* sub-block and variance given by *V*_0_*δα*(1‒*α*) (and is thus uncorrelated across different individuals). Note that in generating offspring trait values, we do not use eq. (1) and hence make no assumptions about linkage equilibrium (except in drawing the random component *ζ*, which goes to zero in the large *L* limit).

This process is repeated in each generation for each descendant individual carrying at least some fragment of the introduced block for some pre-decided number of generations or until no portion of the introduced block survives in the population. At the end of each generation, we ascertain the number of descendants carrying at least some fragment of the introduced block, the total length of introgressed genome they carry, and various moments of the lengths and trait values of the fragments of the introduced block segregating in the population. For each set of parameters, we obtain various statistics by averaging over a large number of realizations (~10^4^) of this process, each corresponding to a different Brownian path.

### 1.3 Modeling the initial spread of the haplotype as a branching process

The above formulation describes a branching process, in that it ignores correlations between the offspring number of different individuals due to effects such as density-dependent regulation, which are required to keep population size fixed. Thus it can only describe short-term introgression into a finite population. Note however that it is a branching process *conditioned* on a particular Brownian path *z*(*x*) with end points at 0 and *y*_0_. Thus the probability *Q_t_*(*y*_1_, *y*_2_) that a genomic block with end-points at *y*_1_ and *y*_2_ has no surviving descendants *t* generations later, must also be conditioned on this Brownian path. We can write the following recursive equation relating the ‘extinction probability’ *Q*_*t*+1_(*y*_1_, *y*_2_) at time *t*+1 to the corresponding probability at time *t*:

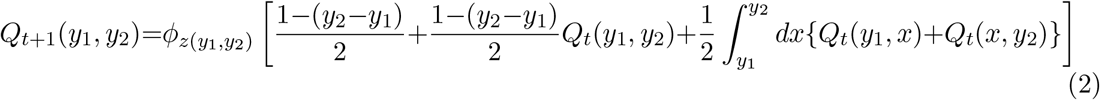

Here *z*(*y*_1_, *y*_2_) is the trait value associated with the genomic region [*y*_1_, *y*_2_], and is specified by the Brownian path. Further, *φ_z_*[*x*] is the moment generating function of the number of offspring of an individual carrying a block fragment with trait value *z*. It is given by *φ_z_*[*x*]=exp(−*w*(*z*)(1−*x*)) under the assumption that individuals have a Poisson-distributed number of offspring with mean *w*(*z*)=2exp(*βz*). Thus *φ_z_*[*x*] is non-linear in *x*, resulting in recursions (eq. (3)) that are also non-linear. Note that φz is technically a functional, since its argument is obtained by integrating over *y*.

The first term in the square brackets represents the probability that the block (of map length *y*_2_−*y*_1_) is not split by recombination and not transmitted to the offspring. The second term describes events in which the entire block is transmitted to the offspring without being split by recombination, but then is completely lost among the descendants of this offspring within the next *t* generations. The third term describes recombination events which split the block, resulting in transmission of a fragment of the block to the descendant. The integral is over the map position *x* of the crossover point. It involves the probability that the immediate offspring inherits the [*y*_1_, *x*] or the [*x*, *y*_2_] fragment, and that this fragment is subsequently lost among the descendants of this offspring within the next *t* generations. The sum of these three terms gives the net probability that there are no surviving descendants of the haplotype after *t* generations in the pedigree branch involving *one* of its offspring. The probability of extinction of the introduced haplotype within two such pedigree branches is the square of this net probability, and so on for three or more branches. This is what allows extinction probabilities to be expressed in terms of the moment generating function of the offspring distribution.

Equation (2) describes the time evolution of extinction probabilities for a particular genomic block described by a particular Brownian path. In order to calculate the extinction probability (or other statistics), averaged over different Brownian paths, all characterized by the same value of V0, we introduce an *approximate* branching process. This approximate branching process is not conditioned on any particular path, but treats the trait values of two or more descendants inheriting portions of the parent block as *independent* random variables. These random variables have a distribution (specified by eq. (1)) which is only conditional on the map length and trait value of the parent block. This approximation thus ignores the correlations in trait values of offspring that inherit overlapping or even complementary tracts of a parent block. The underlying assumption is that these correlations are weak or at the least, do not affect aggregate distributions, e.g., the distribution of the total number of descendants on average.

It follows from the above arguments that under the approximate branching process, the extinction probability of a block depends only on its map length *y* and trait value *z*. Defining *Q_t_*(*y*, *z*) as the probability that a block of map length *y* and trait value *z* has no surviving descendants *t* generations after it enters the population, we can write the following recursive equations for *Q_t_*(*y*, *z*) under the approximate branching process:

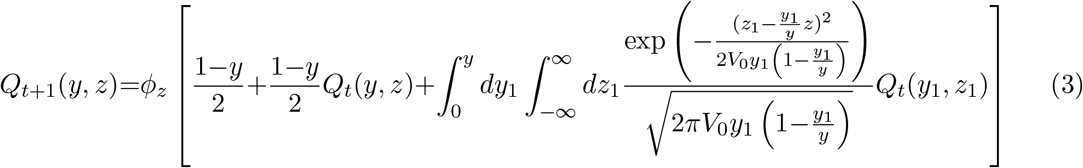

Equation (3) is similar to eq. (2) except that now the third term involves an integral over the map length *y*_1_ and trait value *z*_1_ associated with the descendant block. Note that *z*_1_ is now a random variable drawn from the distribution in eq. (1), and is not pre-specified by the chosen Brownian path, as in eq. (2).

Equation (3) is analogous to eq. 1 in Baird et al. (2003), but unlike that equation, involves extinction probabilities that depend on both the length and the trait value of the introduced block. This reflects the fact that blocks evolve under the joint influence of selection and recombination in our model, in contrast to the neutral dynamics modeled in Baird et al. (2003).

To numerically iterate eq. (3), we discretize (*y*, *z*) space and replace the two-dimensional integral by a double summation. This can introduce some error at long times when the typical length of surviving blocks becomes smaller than the discretization interval. In general, however, we find that predictions of the approximate branching process (as in eq. (3)) show excellent agreement with results of the direct simulations described above.

We can also obtain the Laplace transform of the joint distribution of the number, map lengths and trait values of descendant blocks at time *t*+1 in terms of the corresponding Laplace transform at time *t*, as in Baird et al. (2003). In Appendix S2 of the SI, we show how these can be used to recursively compute moments of the total number of descendant blocks and the total amount of introgressed genome. In particular, we can write down recursions for the expected number *E*[*N_t_*|_*y*_,*z*] of descendants of an introduced block with initial map length *y* and trait value *z* as follows (see Appendix S2):

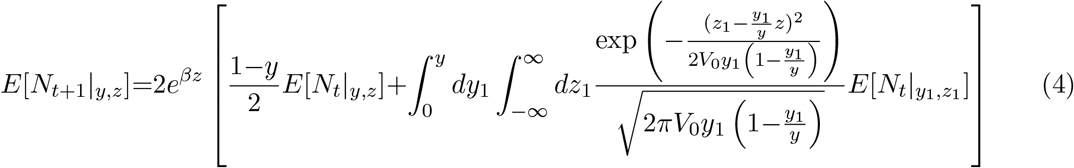

#### Effective parameters governing introgression

Taking the continuous time limit of eq. (4) yields an integro-differential equation for the average number of descendants (see Appendix S3 of the SI). This equation can be written in terms of the map length *y*, the rescaled trait value 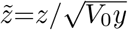, rescaled time *τ*=*yt*, and the ratio of the selection strength to recombination strength, given by 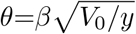:

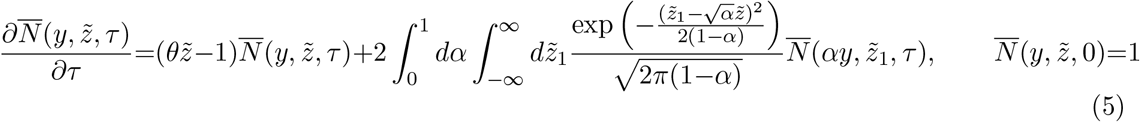

Here 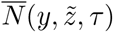 denotes the expected number of descendants of a block of map length *y* and effective trait value 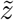, at rescaled time *τ*. Note that the parameters 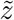 and *τ* (as well as *θ*) are themselves functions of *y*. An identical integro-differential equation describes the evolution of the average total amount of introgressed genetic material 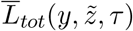, but with the initial condition 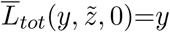. In Appendix S3, we argue that these equations must have solutions of the form 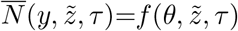 and 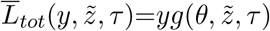. Thus, the total number of descendants depends on the map length of the introduced haplotype only implicitly via the rescaled parameters *θ*, 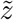, *τ*. Similarly, the genic variance enters into the equations only via *θ* and 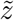.

The above discussion highlights an important difference between the infinitesimal model and the selection model assumed in other studies of multi-locus introgression (e.g. Barton 1983), where trait value is deterministically associated with block length. With deterministic inheritance of trait values, the effective parameter governing introgression is the product 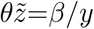, while in the infinitesimal framework, the parameters *θ* and 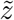 independently influence dynamics (see Appendix S3 for more details).

### 1.4 Individual-based simulations of long-term introgression into a finite population

To determine the time scale over which the above formulation (for introgression into an infinite population) breaks down for a finite population of size *N*, we also perform individual-based simulations of finite populations. These simulations are initialized by randomly choosing a diploid individual in the population to be the carrier of the introduced haplotype. The introduced haplotype is generated by sub-dividing into *L*=2^*R*^ sub-blocks, and iteratively choosing the additive contribution of each sub-block such that the contributions sum to *z*_0_ (as described above). The remaining *N*‒1 individuals are assumed to be identical and have zero genic variance.

In each generation, *N* individuals are created as follows. Two diploid parents are chosen for each offspring by sampling individuals in the previous generation in proportion to their fitness, defined as *e^βz^*. Here *z* is the sum of the trait values associated with the two haplotypes carried by the individual. Subsequently, a gamete is created from each parent either with no recombination (probability 1−*y*_0_) or with a single crossover (probability *y*_0_) between parental haplotypes. In the absence of recombination, one of the two parental haplotypes is chosen with equal probability to be the gamete. With recombination, one of the *L*‒1 junctions between sub-blocks is chosen randomly to be the crossover point; the gamete then inherits all sub-blocks to the right of this junction from one parental haplotype and all sub-blocks to the left of the junction from the second haplotype. The two gametes generated in this way then together form a single diploid individual. All statistics are calculated by averaging over several replicates of the population, each initialized with a different realization of the introduced haplotype.

Note that in the individual-based simulations, we are essentially simulating a genomic block with *L* discrete loci uniformly distributed over a fixed map length. As before, we verify that all quantities of interest are insensitive to the exact value of *L* for large enough *L*, so that this model mimics the infinitesimal model with linkage described above (see Appendix S4 in he SI).

## 2 Results

We first consider the case where a genomic block with trait value *z*_0_=0 enters the native population. This block is *initially* neutral with respect to the native population. However, its immediate descendants inherit fragments associated with non-zero trait values, resulting in a change in the average trait value in response to selection, with the response being stronger for higher values of 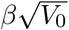. This is in contrast to the purely neutral case (Baird et al. 2003), where there is no variability along the genome (*V*_0_=0) or alternatively no selection (*β*=0), so that descendants always inherit neutral fragments of the originally neutral genome.

The distinction between the 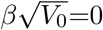 limit considered in Baird et al. (2003) and the 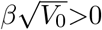 scenarios we study here is less obvious at long time scales. As time progresses, recombination splits the original block into smaller and smaller fragments that contribute less and less to trait value and are thus expected to be effectively neutral. However, selection favours descendants carrying larger fragments with significantly positive contributions to trait value, and thus tends to amplify the frequency of such fragments in the population, in opposition to recombination. Does the resultant *long-term* signature of introgression in the presence of selection differ from the neutral 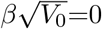 expectation, under the infinitesimal model? This is one of the questions we consider below.

Before computing various statistics of the descendants of the introduced block and exploring their dependence on parameters such as *y*_0_, 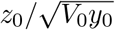 and 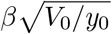, it is useful to visualize a few random realizations of the process (for introgression into an infinite population). Figures 1(a) and 1(c) show two snapshots of the process at *t*=20 and *t*=80 respectively, while figs. 1(b) and 1(d) in the right panel show the analogous snapshots for another realization of the process, corresponding to a different Brownian path. The x-axis denotes physical positions along the genomic region spanned by the introduced block (here chosen to have map length *y*_0_=0.25). The numbers on the y-axis index the descendants of the introduced block, and are different in figs. 1(a) and 1(c) due to the larger number of descendants carrying introgressed material at *t*=80 than *t*=20. Each horizontal line within a plot represents an individual genome. The coloured segment depicts the fragment of the genome that has descended from the introduced block; the colour encodes the trait values associated with the fragment (see accompanying colour scales).

**Figure 1:**
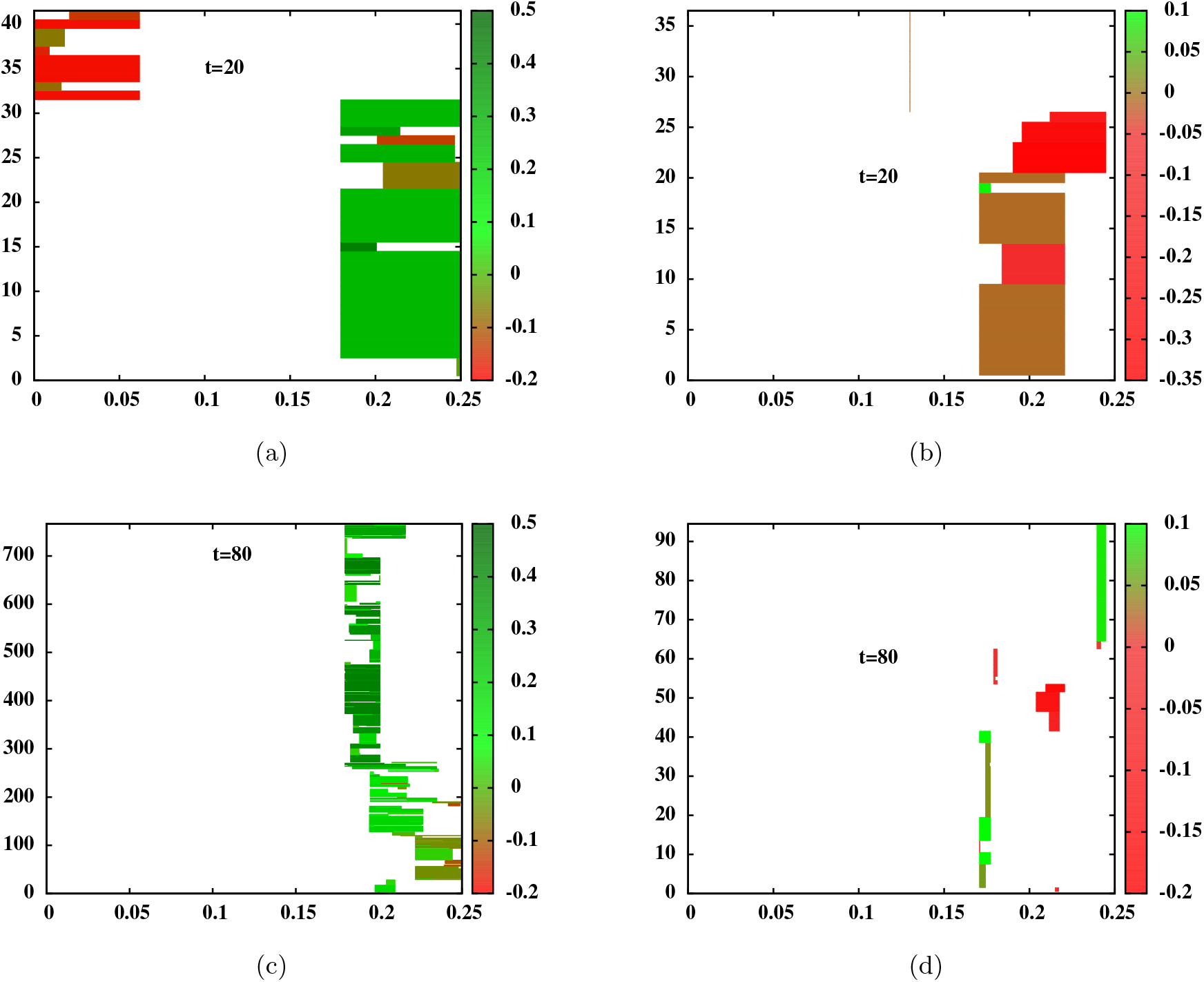
Snapshots of descendants of the introduced genome at *t*=20 (fig. 1(a)) and *t*=80 (fig. 1(c)) for a single realization of the introgression process for an initially neutral block (*z*_0_=0) with map length *y*_0_=0.25 and 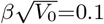. Figures 1(b) and 1(d) show the corresponding snapshots for a second (independent) realization, obtained for introgression of a different genomic block (described by a different Brownian path). Each horizontal line represents the genome of an individual carrying a fragment of the introduced block; the coloured portion represents the introgressed fragment while the white portions represent native blocks. The trait value associated with each introgressed fragment is encoded by the colour of the block (green for positive trait values and red for negative values) and can be read off from the accompanying colour scale. The y-axis of each plot indexes the descendants carrying introgressed block fragments— e.g., in fig. 1(a), there are 41 descendants of the introduced block at *t*=20; thus, there are 41 lines representing 41 genomes in fig. 1(a). The x-axis shows positions along the genomic region influencing trait value (here having map length *y*_0_=0.25). Both realizations are obtained from simulations of introgression into an infinite native population.

In each realization, descendant blocks are longer at *t*=20 than at *t*=80 (on average). Different individuals carry blocks associated with different trait values, with both deleterious (*z*<0 or red) and favourable (*z*>0 or green) blocks present in the population at appreciable frequency at *t*=20. These blocks can themselves be viewed as mosaics of smaller sub-blocks with positive and negative contributions to trait value. As time progresses, recombination tends to isolate small sub-blocks from their genomic backgrounds, allowing selection to amplify the frequency of those sub-blocks that have significantly positive contributions. Thus, at *t*=80, the surviving fragments of the introduced block are mostly associated with positive trait values (fewer red blocks).

Note that although the two Brownian paths corresponding to the two replicates are drawn from the same distribution and hence have similar statistical properties, the two replicates look quite different. For instance, in the first realization (left panel), the introduced genome has a small sub-block with a significantly positive effect embedded within it (as evident in the large number of dark green segments between genomic positions 0.179 and 0.216 in fig. 1(c))— this results in rapid introgression, with more than 700 individuals carrying introgressed genetic material at *t*=80. In the second realization (right panel), the original genome contains moderately favourable sub-blocks; as a result, the spread of these sub-blocks is more modest (~90 descendants at *t*=80). Interestingly, surviving blocks appear to be longer in the first realization than in the second (e.g., at *t*=80), which suggests a correlation between the trait values and lengths of surviving fragments.

In Appendix S5 of the SI, we also consider replicate simulations corresponding to the *same* introduced genome (represented by the same Brownian path), which however still differ from each other in their stochastic histories of birth and recombination events. These replicates are quite variable as well (see fig. 3 in Appendix S5, SI), suggesting that the stochasticity inherent in reproduction and recombination may also be an important source of variability between replicates, especially in the early phases of introgression.

To further clarify these effects, we compute various statistics associated with individual fragments of the introduced block, by first calculating the mean (or variance) of block lengths and trait values for any one realization of the introgression process, and then averaging the mean (or variance) over many such realizations (each corresponding to a different realization of the Brownian path). The average trait value of surviving fragments is positive and increases with time (fig. 2(b)) even as the length of fragments decreases (fig. 2(a)). This is consistent with fig. 1 and the observation that a large neutral block typically has small positively selected fragments embedded within it, which can be dislodged by a few generations of recombination. The increase in average trait values is thus most dramatic for large values of 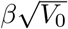, i.e., when there is high genetic variability along the introduced genome (which makes it more probable that it has constituent sub-blocks with significantly positive fitness effects) and if there is strong selection (which amplifies the frequency of such blocks).

**Figure 2:**
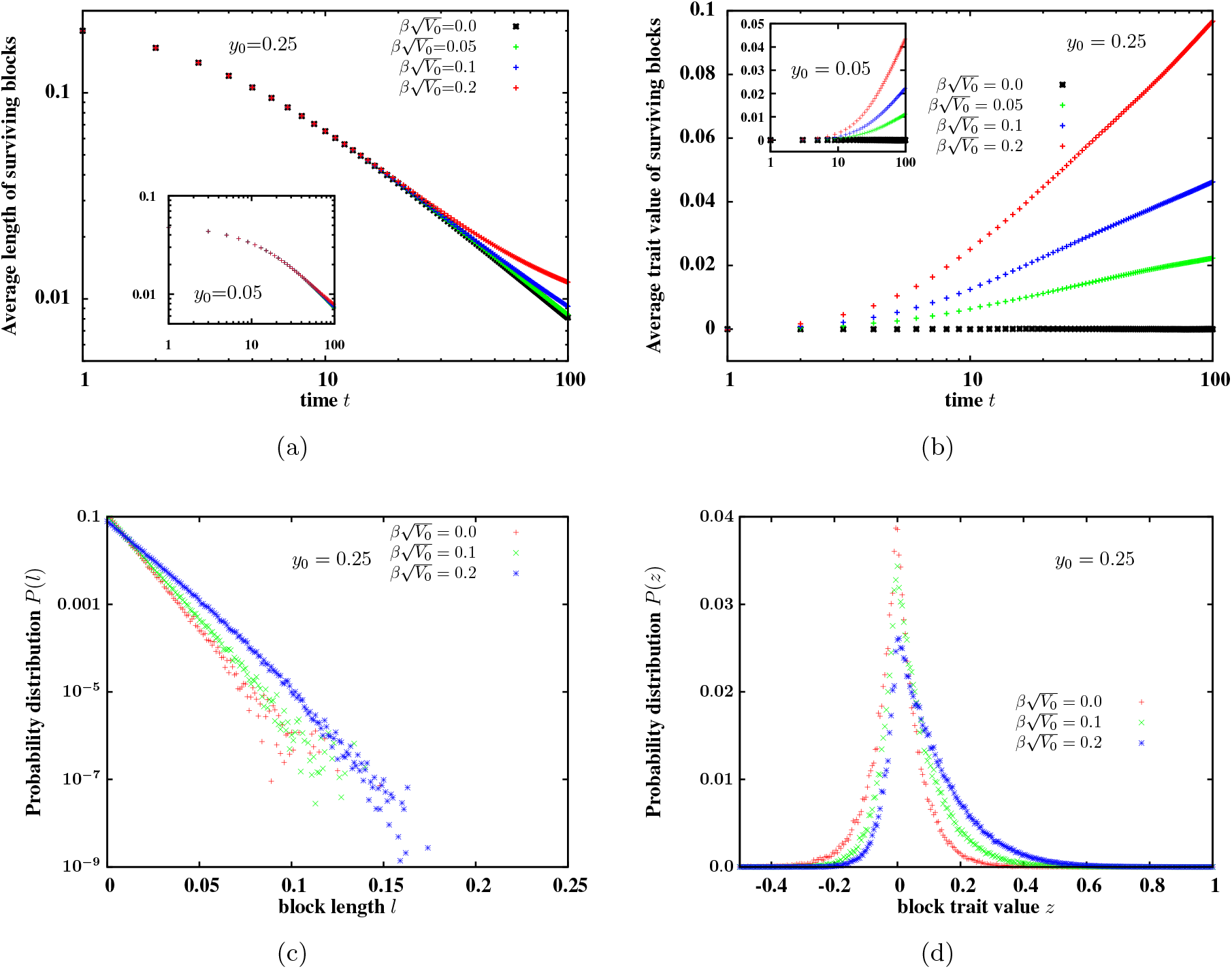
(a) Average length of surviving fragments of the introduced block in a population with at least one such fragment, and (b) average trait values of surviving fragments, versus *t*, the number of generations since the initial hybridisation event. Main plots show statistics for original block length *y*_0_=0.25 and inset for *y_0_*=0.05. (c) Probability *P*(*l*) of detecting an introgressed fragment of map length *l*, *t*=100 generations after the initial hybridisation event, for various values of 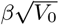. (d) Probability *P*(*z*) that an introgressed fragment present in the population at *t*=100 is associated with trait value *z*, for various 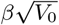. The map length of the selected region is *y*_0_=0.25 and the trait value associated with the initial introduced block is *z*_0_=0 in all plots. All plots are obtained from simulations of the introgression process into an infinite native population, by averaging over 10^4^ realizations of the process.

However, once a small positively selected genomic sub-block has been isolated, further splitting dilutes its selective advantage. The fact that average trait value associated with descendant fragments continues to increase over several hundred generations (in spite of recombination), is due to selection, which favours individuals inheriting the whole sub-block over descendants inheriting smaller portions of the sub-block. To see this, note that a sub-block with map length *y*_*_ and trait value *z*_*_ is passed on as it is (i.e., without splitting) to an average of *w*(*z*_*_)(1‒*y*_*_)/2 descendants. Thus any sub-block with *w*(*z*_*_)(1‒*y*_*_)/2>1 spreads exponentially through the population as a *non-recombining unit,* and dominates various statistics such as those in figs. 2(a) and 2(b). The long-term increase in average trait value among descendant blocks is thus explained by the increase in frequency of blocks with *w*(*z*_*_)(1−*y*_*_)/2>1.

This also implies that when selection is strong, fairly large sub-blocks with moderately positive contributions (*y*_*_<1‒e^−*β*_*z*_*__^) can spread exponentially (without being split). As a result, the average length of surviving fragments decays significantly slower than the neutral expectation for large values of 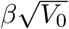 (fig. 2(b)). Concomitantly, the distribution of block lengths *l* is shifted towards large *l* (fig. 2(c)) while the distribution of trait values among surviving blocks is significantly skewed towards positive values (fig. 2(d)) when selection is strong 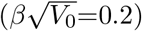.

Signatures of selection are evident in other statistics associated with the population as a whole (fig. 3). Figure 3(a) shows the survival probability *P_surv_*(*t*) of at least some part of the introduced block (i.e. the probability that one or more fragments of this block survive in the population *t* generations after the original hybridization event) as a function of *t*, for various values of 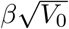. Figure 3(b) shows the average number 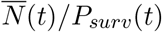 of descendants carrying fragments of the introduced block, conditional on the survival of at least one such descendant. Figure 3(c) depicts the total amount 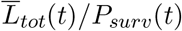 of introgressed genetic material (obtained by summing over the map lengths of all fragments of the introduced block present in the population), again conditional on survival of some part of the block. The main plots and insets show results for blocks of map length 0.25 and 0.05 respectively. The points are obtained from simulations of introgression into an infinite population, while lines show the predictions of the approximate branching process. Note the excellent agreement between the two, at least at short times.

**Figure 3:**
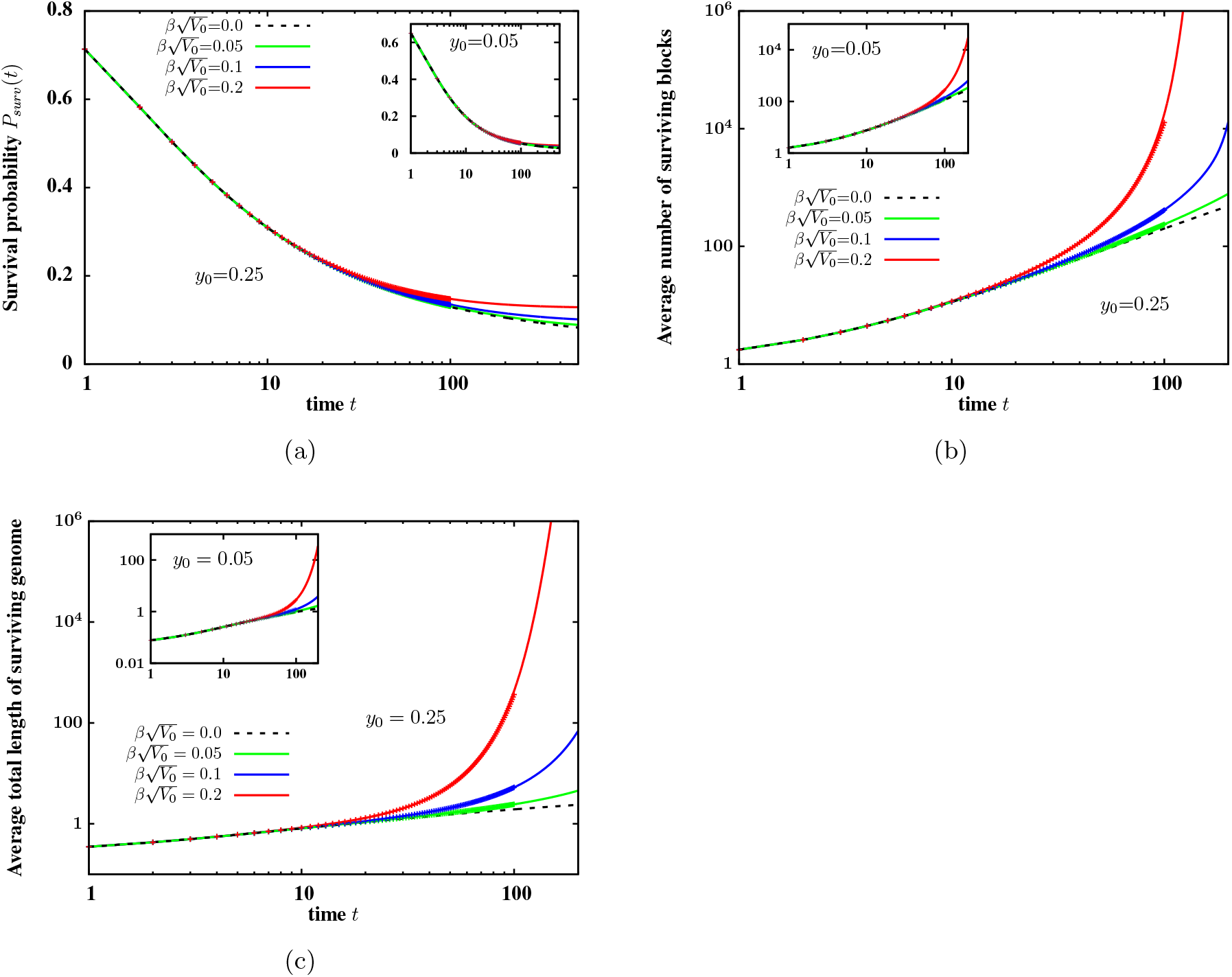
(a) Survival probability of at least one descendant of the initially neutral (*z*_0_=0) introduced block (b) average number of descendants of the block (conditional on survival of some part of the block) and (c) average of the total map length of introgressed genetic material in the population, versus time *t* after the initial hybridization event. The averages in plots (b)-(c) are normalized by the survival probability at time *t*. Points are calculated from simulations of the introgression process into an infinite native population, while lines show predictions of the branching process approximation and are obtained from numerical iteration of eqs. (3) and (4). Both inset and main plot depict statistics for 4 different values of 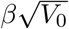. Main plots show statistics for original block length *y*_0_=0.25 and inset for *y*_0_=0.05. Deviations from the neutral expectation (dashed line) become evident only after several tens of generations.

The probability *P_surv_*(*t*) follows the neutral expectation (dashed line) over ~30 generations irrespective of the strength of selection, but decays more slowly for larger values of 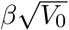 over longer time scales. Marked deviations from neutral dynamics can also be observed for 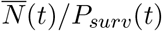 and 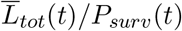 (figs. 3(b) and 3(c)). At long times, these grow *exponentially,* as opposed to the linear in time spread of introgressed material predicted in the neutral case by Baird et al. (2003) (see also dashed black lines in figs. 3(b) and 3(c)). This is consistent with our earlier observation that at long times, surviving fragments of the block must have positive trait values. Directional selection then results in exponentially fast introgression of some of these fragments (essentially as non-recombining units).

Another noteworthy feature of fig. 3 is the increased likelihood (fig. 3(a)) and higher rate (figs. 3(b)-3(c)) of introgression, when the map length *y*_0_ of the introduced block is large (compare inset and main plot for any value of 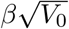. Longer genomic blocks have a twofold advantage— the overall probability that a genomic block of length *y* is passed on (either in part or wholly) to a haploid gamete increases as (1+*y*)/2 (for *y*≪1), resulting in a *transmission* advantage for longer blocks; moreover, longer blocks display higher variability (proportional to *V*_0_*y*) and are thus more likely to contain small beneficial fragments, resulting in a long-term selective advantage for their descendants.

### Introgression into a finite population

Deviations from the neutral expectation typically manifest several generations (~30 generations in fig. 3) after the initial hybridisation event. However, the analysis described above (for introgression into an *infinite* native population) is not expected to be quantitatively accurate for finite populations of size *N* at very long time scales. Moreover, in the infinite population case, long-term dynamics of trait values and block lengths is predicated on the indefinite exponential spread of small blocks with significantly positive trait contributions (*w*(*z*_*_)(1‒*y*_*_)/2>1), which is again unrealistic for any finite *N*. To determine the regime of validity of the above analysis, we compare various statistics for finite *N* (obtained from individual-based simulations) with infinite population predictions.

Figure 4(a) shows the average total amount of surviving introgressed genome 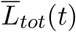 in the population as a function of time *t* after the hybridisation event. The infinite population model (lines) agrees with finite *N* predictions (points) over a short time scale, which increases logarithmically with *N* (for 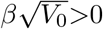). This is explained by the fact that the average number of descendants (as well as 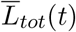) grows exponentially in time in an infinite population (figs. 3(b) and 3(c)), and thus becomes comparable to any fixed population size *N* at a time that scales as log(*N*). Further, *L*_*tot*_(*t*) and the number of descendants have long-tailed distributions (under the infinite population model)— for example, the expected value of the number of descendants of an initially neutral block in the presence of strong selection (*z*_0_=0, *y*_0_=0.25, 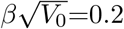) is ~10000 at *t*=100, while the standard deviation is 70000. Thus, the typical (as opposed to the average) number of descendants approaches any fixed *N* even more rapidly.

**Figure 4:**
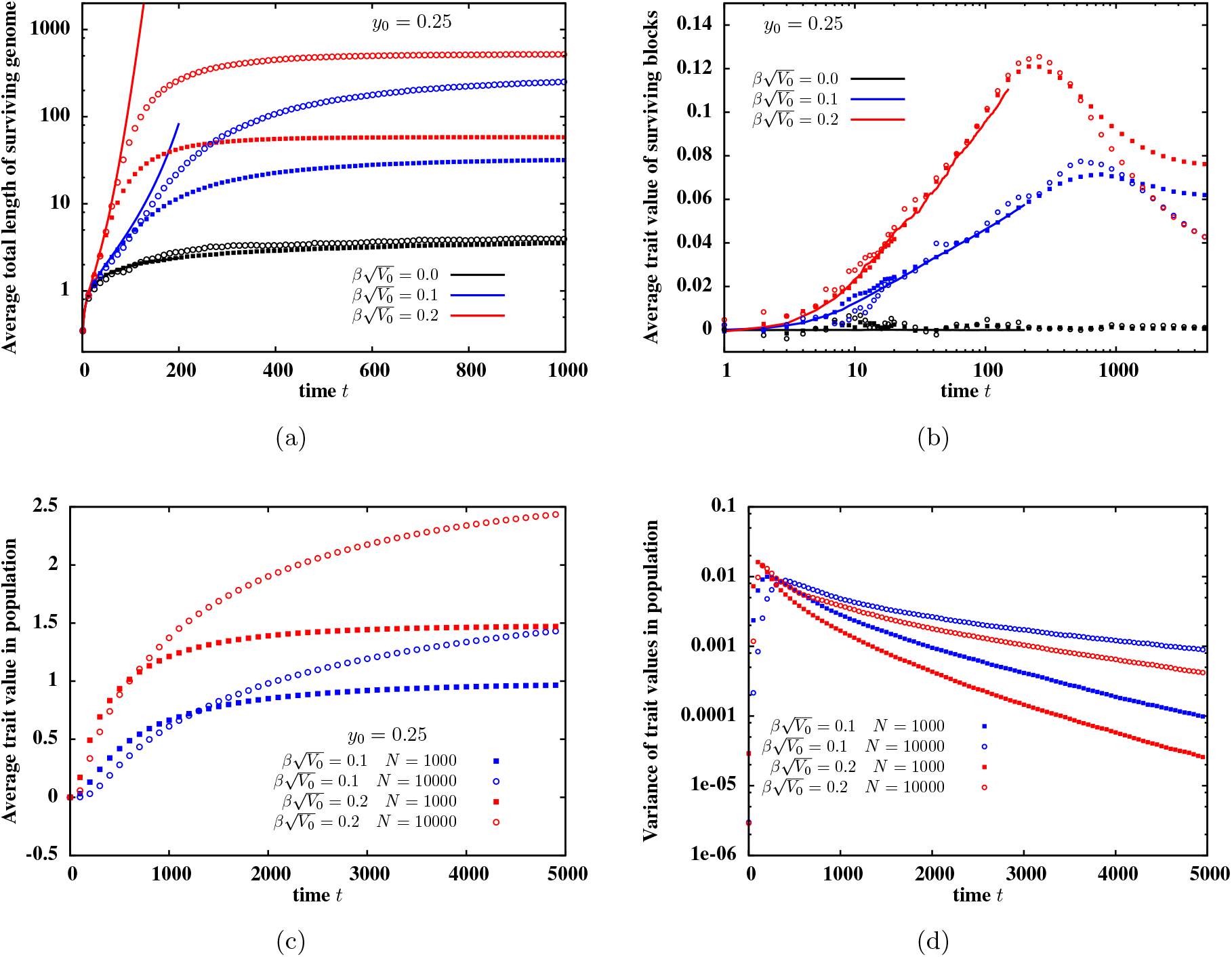
(a) Total amount (length) of introgressed genome surviving in the population *t* generations after the initial hybridisation event as a function of *t*, for various values of 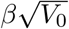. (b) Average trait value associated with a surviving block, versus *t*. (c) Average trait value associated with individuals in the population, versus *t*. (d) Variance of trait values associated with individuals in the population (averaged over replicates), versus *t*. Lines in (a)-(b) represent predictions of the infinite population model while points are from individual-based simulations of *N*=10^3^ (squares) and *N*=10^4^ (circles) populations. All simulations are with *L*=2^12^ discrete loci. The infinite population predictions match statistics calculated for finite populations over an initial time scale that increases weakly with *N*.

Once the number of descendants of the introduced block becomes comparable to the population size, mating between individuals carrying the introgressed genome becomes frequent, resulting in offspring who inherit *multiple* introgressed fragments. For instance, with strong selection 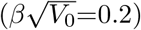, the average number of introgressed fragments per individual ranges from ~5.5 (for *N*=1000) to ~3 (for *N*=100000) as early as at *t*=200.

This has interesting implications for the evolution of trait values and hence, the rate of adaptation in finite populations (as shown in figs. 4(b) and 4(c)). The average trait value associated with descendant blocks in a finite population follows the infinite population prediction over a short time scale (that becomes larger with increasing *N*), but then starts declining over longer times (fig. 4(b)). This is explained by the fact that a short block with a significantly positive trait contribution ((1‒*y*) exp(*βz*)>1) cannot keep spreading exponentially through a finite population. This exponential spread was responsible for the continued increase in trait values of surviving blocks within the infinite population (see above).

However, since a typical descendant in a finite population accumulates multiple introgressed blocks at long times, the average trait value associated with *individuals* keeps increasing (fig. 4(c)) even as the trait value of individual blocks falls. The average trait value initially increases faster in the smaller (*N*=10^3^) population in which mating between descendants of the introduced block starts occurring soon after the hybridisation event (fig. 4(c)). However, the fixation of subblocks and the accompanying fall in genetic variability is also faster in the *N*=10^3^ population (fig. 4(d)). The higher genetic variance in the *N*=10^4^ population allows for a longer selection response and a greater net advance in the trait value (fig. 4(c)).

Thus, in the present model, adaptive introgression into a finite population involves two distinct processes occurring on different time scales. The first phase is characterised by splitting of the original block by recombination, the separation of genomic fragments with positive fitness effects from their deleterious background and the amplification of the frequency of such fragments by selection. During this phase, the average trait value among descendants carrying introgressed genetic material increases logarithmically in time and is well predicted by the infinite population model. The second phase is characterised by increased probability of mating between individuals carrying introgressed genetic material and the emergence of individuals who bear multiple introgressed fragments (with positive contributions to trait value), and have a strong selective advantage. This causes a more rapid increase in average trait value in the population at longer time scales, which cannot be predicted by the infinite population model.

Nevertheless, the infinite population model provides valuable intuition about the initial phases of introgression, and is easier to simulate since it requires us to only keep track of introgressing block fragments. We now use the infinite population framework to explore how blocks that are originally non-neutral (with trait value *z*_0_≠0) spread through the native population during the early phases of introgression.

### 2.1 Favourable blocks

Consider a scenario where a single copy of a favourable genomic block (with 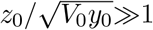) enters the native population. Unlike in the case of a selectively neutral (*z*_0_=0) introduced block, here we expect the trait values associated with descendants to become less positive from generation to generation as the original block splits up into smaller fragments. This leads to the question: is a single positively selected locus (corresponding to the *y*_0_→0 limit) more likely to introgress into a homogeneous native population than a long genomic block with the same selective advantage? We first consider the case of introgression into an infinite native population. As before, we follow various attributes such as *P_surv_*(*t*), 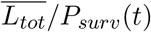 and the average size and effect of descendant blocks through time, but now with a focus on how these vary with *y*_0_ (the map length of the introduced block).

Figure 5(a) shows *P_surv_*(*t*) versus t for various values of *y*_0_ for two different strengths of selection (main plot and inset). For both values of 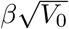, the survival probability is minimum for the *y*_0_→0 case (corresponding to a single discrete locus) and increases with *y*_0_, but more weakly than in the case of the neutral block. The weak dependence of *P_surv_*(*t*) on *y*_0_ reflects the tension between two opposing effects— the higher (overall) probability of transmission of longer blocks to the gamete during meiosis versus the fact that long blocks are more likely to be split by recombination and hence transmit only a part of their selective advantage to the next generation.

**Figure 5:**
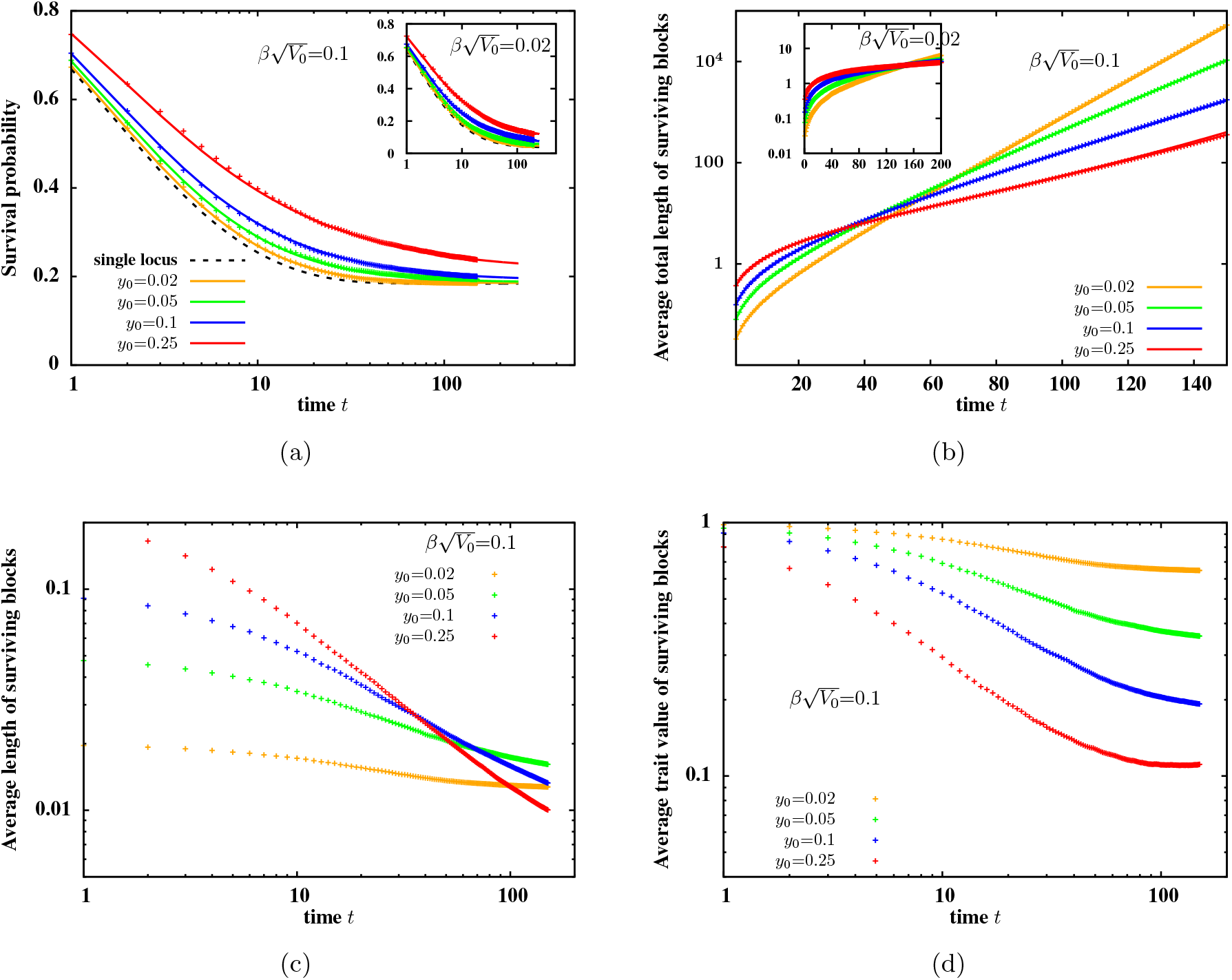
(a) Survival probability of at least one descendant of an originally beneficial 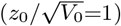 introduced block (b) average of the total map length of introgressed genetic material in the population (conditional on survival of some part of the introduced block) (c) average length of surviving blocks and (d) average trait values of surviving block, versus *t*, the number of generations since the initial hybridisation event. Points are calculated from simulations of the introgression process into an infinite native population, while lines in (a)-(b) show predictions of the branching process approximation and are obtained from numerical iteration of eqs. (3) and (4). Both inset and main plot depict statistics for 4 different values of 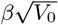. Main plots show statistics for original block length *y*_0_=0.25 and inset for *y*_0_=0.05.

Within surviving lineages, blocks that are originally short leave more descendants and result in a higher total length of introgressed genome in the population (at long times) than long blocks with the same selective advantage (fig. 5(b)). This is due to the fact that short blocks undergo fewer divisions and less dilution of selective effect. This is also evident in fig. 5(d) which shows that the long-term average trait value associated with surviving fragments is higher when the introduced block is small.

Since long blocks break up faster than short ones, a hundred generations after the hybridisation event, fragments of the *y*_0_=0.25 genomic block are already shorter on average than fragments of an originally smaller (e.g., *y*_0_=0.02) block that had the same trait value (fig. 5(c)). Thus, at long times, the average length of descendant blocks depends non-monotonically on the map length of the introduced block— surviving fragments are longest when the initial selective advantage is spread over a genomic block of *intermediate* length (fig. 5(c)).

As in the *z*_0_=0 case, the infinite *N* model fails to describe long-term introgression into finite populations. In a finite population, the average trait value associated with descendants individuals actually increases at long time scales, even as trait values of descendant blocks falls (results not shown). As before, this is due to the emergence of individuals carrying multiple introgressed fragments. Thus, recombination can be thought of as playing a dual role during the introgression of a beneficial genomic block. In the initial phase, recombination splits the original block into smaller and smaller fragments, thus diluting the selective advantage of descendants carrying block fragments over time. In the later phases, recombination re-assembles those fragments that have survived selection (i.e., have sufficiently positive contributions), resulting in genomes with very large positive trait values.

Note that the dependence of quantities such as 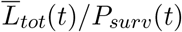 and 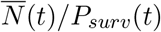 on the initial map length *y*_0_ is qualitatively different for introduced blocks with strongly positive 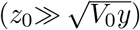 trait values and blocks that are initially neutral (*z*_0_~0), suggesting that there could be a crossover between these two kinds of dependence at some critical value of *z*_0_. To investigate this, we plot the average number of surviving descendants 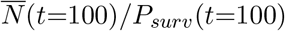 a hundred generations after the initial hybridisation event, as a function of map length *y*_0_ of the selected genomic block, for various initial trait values *z*_0_ (fig. 6). For low or zero *z*_0_, the number of descendants increases with increasing *y*_0_. This is explained by the higher genetic variation associated with longer blocks. When the introduced block is neutral (or nearly neutral) with respect to the native population, the response to selection must be driven by the release of this variation by recombination. By contrast, when the initial trait value *z*_0_ is high, initial introgression depends more on the selective advantage of the introduced block over native blocks, and less on the generation of new variation. In this case, varying *y*_0_ essentially alters the balance between selection and recombination, with longer blocks splitting faster, resulting in a rapid dilution of their selective effect. Thus, in this case, 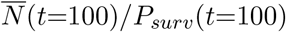 actually falls with *y*_0_. Interestingly, at intermediate values of *z*_0_, the average number of surviving descendants exhibits a *non-monotonic* dependence on y_0_ (see *z*_0_=0.4 and *z*_0_=0.6 curves in fig. 6), signifying a switch from 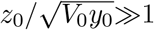 behaviour (associated with strongly advantageous genomic blocks) to 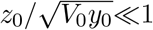 behaviour (associated with initially neutral blocks).

**Figure 6:**
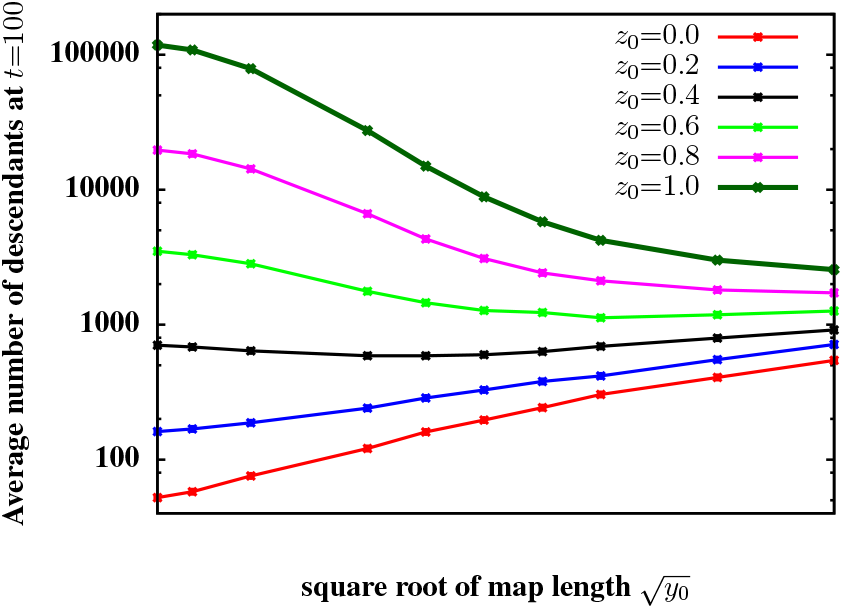
The average number of surviving descendants carrying some part of an introduced block for 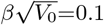, 100 generations after the initial hybridisation event (conditional on survival of at least one such descendant), versus 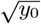, where *y*_0_ is the map length of the introduced block. The different curves correspond to different values of *z*_0_, the (unscaled) trait value associated with the introduced block.

### 2.2 Deleterious blocks

We finally consider the case of a block with *z*_0_<0, which is at a selective disadvantage with respect to the native population. The role of selection in this scenario is somewhat subtle— on the one hand, stronger selection makes it likely that lineages containing (deleterious) fragments of the original block die out in the first few generations itself; on the other, small fragments of the block with even mildly positive contributions to trait value, once separated from the deleterious background, have a strong selective advantage which causes them to take over the population. Thus, while the survival probability (that at least part of the block survives) declines more rapidly for larger 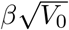 in the first few generations (fig. 7(a)), it also approaches its asymptotic value faster.

**Figure 7:**
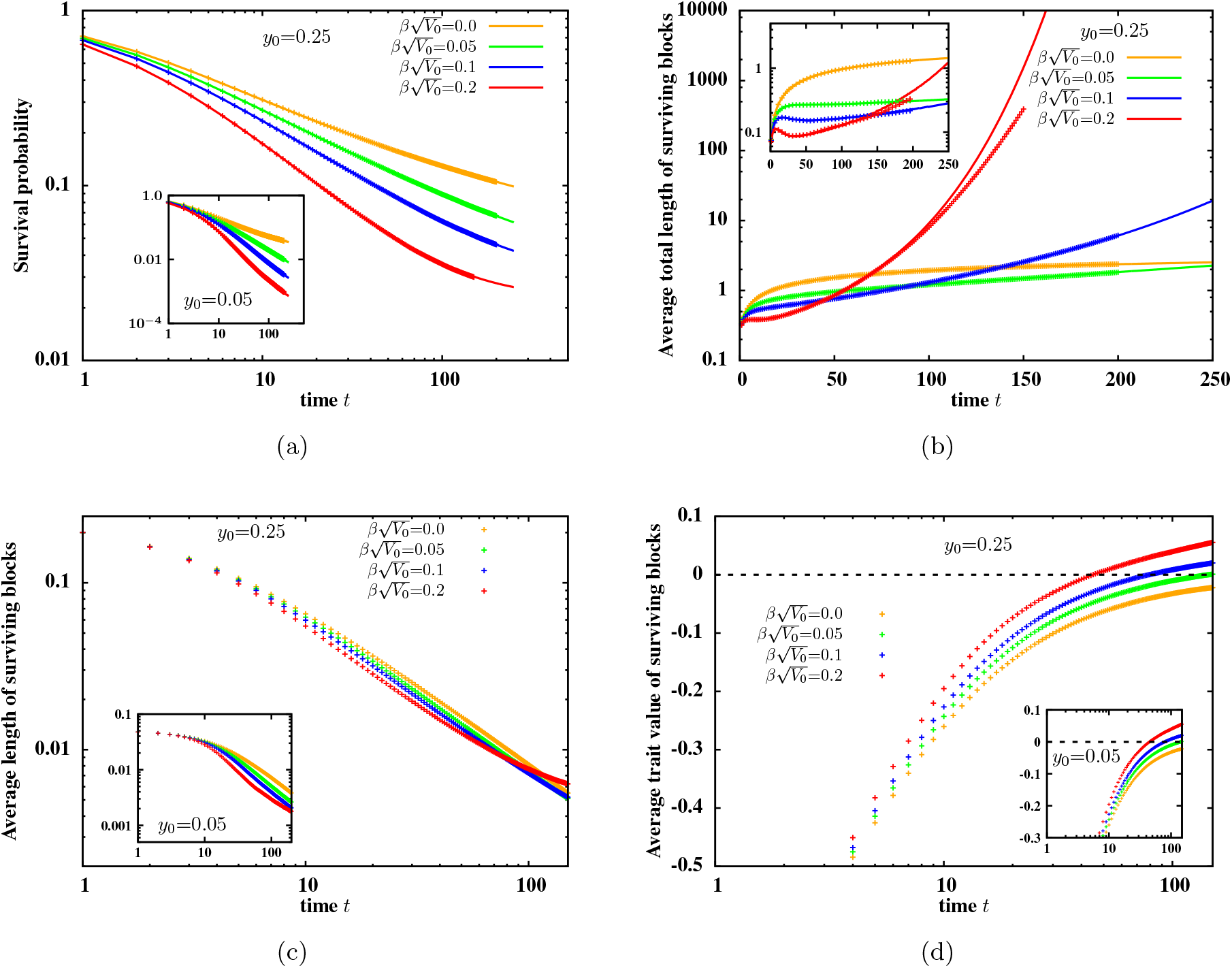
(a) Survival probability of at least one descendant of an originally deleterious 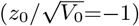 introduced block (b) average of the total map length of introgressed genetic material in the population (conditional on survival of some part of the introduced block) (c) average length of surviving blocks and (d) average trait values of surviving block, versus *t*, the number of generations since the initial hybridisation event. Points are calculated from simulations of the introgression process into an infinite native population, while lines in (a)-(b) show predictions of the branching process approximation and are obtained from numerical iteration of eqs. (3) and (4). Both inset and main plot depict statistics for 4 different values of 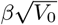. Main plots show statistics for original block length *y*_0_=0.25 and inset for *y*_0_=0.05.

Moreover, the average number of descendant blocks (conditional on survival) and the total amount of introgressed genome they encompass, grows faster for larger 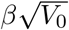, i.e., for stronger selection against the original deleterious block and/or higher genic variance of the block (fig. 7(b)). The trait value associated with surviving descendant blocks also becomes rapidly positive when 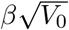 is large (fig. 7(d)). This is consistent with the more general expectation that under strong selective pressures, maladapted lineages must undergo rapid adaptation or are soon lost from the population.

The average length of blocks descended from the originally deleterious haplotype is shorter than that of descendant blocks of the neutral 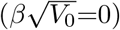 haplotype (fig. 7(c)). Shorter blocks are associated with less negative contributions to trait value (on average) and are favoured by selection— thus, the stronger the strength of selection, the more markedly the average block size deviates from the neutral expectation, at least at short times. At long times, however, the decay in average block size becomes slower, as small positively selected fragments emerge from the deleterious blocks. As before, this can be explained by the fact that selection will oppose further splitting of a positively selected small genomic block and favour descendants who inherit the whole (as opposed to a portion of) this block.

Figures 7(a)-7(b) also show that large deleterious blocks are more likely to survive and also leave more descendants within surviving lineages than small deleterious blocks (insets vs. main plots). This is both because of the higher transmission advantage of large blocks during meiosis as well as the higher segregation variation and superior adaptive potential associated with them.

## 3 Discussion

Our study highlights how the introgression of a block of genome under positive (directional) selection differs qualitatively from neutral introgression, even under an infinitesimal model. Positive selection causes the average number of descendants carrying introgressed fragments, as well as the average total amount of introgressed genetic material in the population, to grow exponentially in time, at least in the initial phase of introgression (while the number of descendants is smaller than the population size). Exponentially fast introgression sets in almost immediately after the initial hybridisation event, in the case of an introduced block with significant selective advantage (fig. 5(b)). For introduced blocks that are neutral or nearly neutral, deviations from the linear (neutral) expectation become evident only several tens of generations after the hybridisation event (figs. 3(b) and 3(c)). This is contrary to the intuitive expectation that under the infinitesimal model, multiple rounds of recombination should progressively dilute the selective effect of descendant blocks, leading to introgression patterns that are essentially indistinguishable from those of neutral introgression over longer time scales. Here, we show that this dilution of selective effect is countered by selection, which tends to amplify moderately sized fragments having significantly positive additive contributions, resulting in their proliferation in the population, essentially as non-recombining units.

Our analysis also suggests that introgressed genomic fragments surviving at long times have an average length which depends on relatively few parameters, such as the trait value and map length of the introduced block, and the strength of selection in the population. For an introduced block with a significant selective advantage, descendant blocks are longest (on average) when the initial selective effect is distributed over a block of intermediate length (fig. 5(c)). In general, the length of surviving blocks is shaped by two opposing effects— the block must be small enough that it is transmitted intact (without recombination) to the majority of offspring, but large enough that it contributes substantially to trait value.

This leads to the question— do genomic signatures associated with the spread of such medium-sized blocks containing many infinitesimally selected loci differ from introgression signatures of a single strongly selected locus? To get a sense of this, we do individual-based simulations in which we also track haplotype diversity *H*, measured as 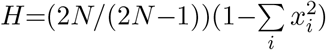, where *N* is the population size and *x_i_* is the frequency of haplotype *i* in the population (Nei and Tajima 1981). The native population is assumed to have 2*N* distinct haplotypes before the hybridization event. We then measure how haplotype diversity changes with increasing map distance from a selected locus or block, as it introgresses and sweeps to fixation (fig. 8). The discrete locus and the extended blocks are associated with the same initial trait value (*z*_0_=1) and experience the same selective pressures (*β*=0.1). Regions of reduced diversity can be observed near the discrete locus (figs. 8(a),8(b)) as well as the short block (figs. 8(c),8(d)), and also near introgressing fragments of the large selected block (fig. 8(e),8(f)). These simulations thus suggest that commonly used diagnostics, based on reductions in linked neutral diversity, may fail to distinguish between the sweep of a discrete locus and the sweeps of short or moderately sized genomic blocks with a large number of infinitesimally selected loci. Note, however, that the hard sweep-like patterns that we observe in fig. 8 may be altered if introgression involves continual migration or the native population is genetically heterogeneous. These issues will be explored in detail in a future study.

Adaptive introgression can be driven either by the introduction of very fit haplotypes into a population or by the generation of new variation via recombination— even when the introduced haplotype is initially neutral or deleterious with respect to the native population. Linkage plays a qualitatively different role during these two modes of introgression (fig. 6). When the introduced haplotype is fit, tight linkage ensures that it is essentially passed on as a non-recombining unit, leading to faster introgression by smaller blocks. When the introduced haplotype is nearly neutral, larger map lengths (weaker linkage) are correlated with more hidden variation and a higher probability that a small positively selected sub-block is embedded within the neutral block. This sub-block can be separated from the neutral or deleterious genomic background by recombination, resulting in a strong response to selection. Thus, in the case of neutral or nearly neutral introduced blocks, introgression is faster for larger map lengths.

It is also worth noting that recombination can play qualitatively different roles during different phases of introgression into a finite native population. When the introduced haplotype is strongly beneficial, recombination dilutes the selective advantage associated with descendant blocks in early stages of introgression (thus acting counter to selection). In later phases, recombination tends to bring together small positively selected sub-blocks (that have survived selection), generating descendant individuals with multiple introgressed fragments and a strong selective advantage. Thus, over longer time scales, recombination accelerates introgression by countering Hill-Robertson interference (Hill and Robertson 1966), and thus crucially determines the net advance under selection.

When the trait under selection is determined by an infinite number of unlinked loci (as in the standard infinitesimal model), the net advance under selection scales with the size *N* of the population (Robertson 1960), and is expected to be infinite for an infinite population. In our simulations of the infinitesimal model with linkage, we observe a much weaker dependence on *N* (see fig. 4(c), where a ten-fold increase in population size causes the net advance to increase only by a factor of 2–3). A related effect was noted by (Weissman and Barton 2012)— they found that even with a constant, high rate of beneficial mutations, the rate of adaptive substitution approaches a limiting value that is independent of population size and mutation rate, but is only constrained by the map length of the mutational target.

An important caveat is that most of the trends described above are only evident for the *averages* of various quantities, where the averaging is over thousands of replicates, each involving a different realization of the introduced block. Individual replicates can however differ dramatically from one another (see fig. 1 above and fig. 3 in Appendix S5, SI) and also from the average. In fact, quantities such as the number of surviving descendants or the total length of introgressed genome have a long-tailed distribution in an infinite population or even in finite populations at short times, such that the standard deviation of these quantities is usually much larger than their mean. Such variability between replicates presents a severe challenge for developing methods that infer population genetic parameters (typically from a single realization of the introgression process, as in fig. 8).

**Figure 8:**
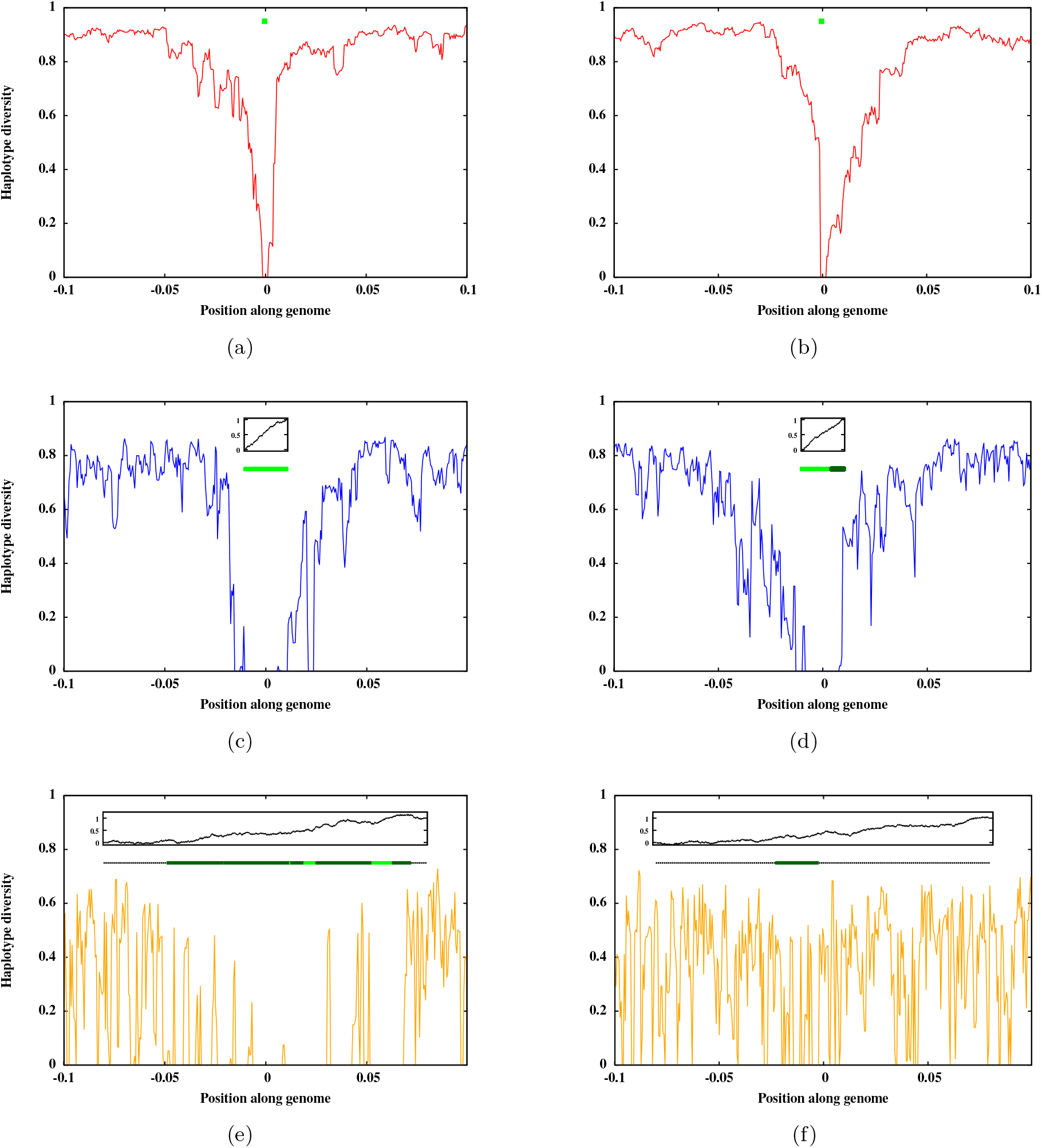
Haplotype diversity along the genome in the native population, a number of generations after the initial hybridisation event in which a single copy of a positively selected locus (figs. (a)-(b)), or a small selected block with map length *y*_0_=0.02 (figs. (c)-(d)), or a larger selected block with *y*_0_=0.16 (figs. (e)-(f)) is introduced into the population. The x-axis represents position along the genome, with *x*=0 denoting the position of the selected locus (in figs. (a)-(b)) or the mid-point of the selected block (in figs. (c)-(f)). The y-axis represents haplotype diversity (defined in text). The light green bars in each figure show those fragments of the introduced block that have been fixed in the population; the dotted segments represent the fragments that have been lost, while the dark green segments represent those fragments that are segregating at some appreciable frequency in the population. The black curves in the insets in figs. (b)-(f) show how the trait value varies along the selected block (with increasing distance from the edge of the block). The two sub-figures in each row depict results from replicate simulations. The trait value associated with the selected locus or block is *z*_0_=1 in each plot; other parameters are *β*=0.1 and 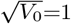. Each figure is obtained from individual-based simulations of populations with size *N*=1000. Haplotype diversity is measured *t*=200 generations after the initial hybridisation event for the case of the discrete locus (figs. (a)-(b)), *t*=500 generations after the initial event in case of the short block (figs. (c)-(d)), and *t*=2000 generations after the initial event in case of the extended block (figs. (e)-(f)).

The high variability in the outcomes of introgression among different replicates reflects two different kinds of underlying stochasticity. First, individual Brownian paths describing the introduced block can be quite different from each other, even if they are drawn from the same distribution. Positive-effect loci may be physically clustered along the introduced block by chance even if there is no population-wide LD in the source population from which the block originated. Rapid introgression ensues when the introduced block happens to contain a genomic tract with strong positive effect (for instance in figs. 1(a) and 1(c)), while introgression is much slower when the clustering is weaker and the constituent tracts only moderately beneficial (figs. 1(b) and 1(d)). This also suggests that introgression would be more unlikely if the source population were under very strong stabilizing selection (which would create negative LD and reduce the likelihood that a genome contains stretches with significant positive effect).

Different replicates corresponding to the same Brownian path can also be quite different from each other (see fig. 3 of Appendix S5 in the SI). This is due to the stochasticity inherent in reproduction and recombination events. Various stretches of an introduced block may be lost from the population just by chance— this is especially true in the initial phases of introgression of a long, nearly neutral block, when segregating sub-blocks are also long and associated with weakly non-zero effects. At longer times, once moderately sized blocks with a substantial contribution to trait values (i.e., *e*^*βz*^(1‒*y*) significantly greater than 1) have been isolated, we expect introgression patterns to be essentially dominated by the proliferation of the sub-block with the largest *e*^*βz*^(1‒*y*. Understanding the role of stochasticity during different phases of introgression and analyzing whether a deterministic model can describe the later phases of introgression, remains an interesting direction for future work.

Most of our analysis is based on simulations of a finite native population or an effectively infinite population (in which the descendants of the introduced block can be thought of as generating a branching tree). However, we find that many statistics obtained from these simulations are actually in excellent quantitative agreement with the predictions of a branching process approximation that neglects correlations in trait value among different descendants of an individual, and also employs eq. (1) which is only true when there is no LD. Negative LD may arise, however, in a finite population under directional selection due to Hill-Robertson interference (Hill and Robertson 1966). It would be interesting to explore whether an economical description for inheritance within the infinitesimal framework, analogous to eq. (1), can be found in the presence of linkage and linkage disequilibrium (as generated by Hill-Robertson interference or stabilizing selection).

Note that while the infinitesimal model strictly assumes an infinite number of selected loci (in this case within a limited genomic region), in practice, its predictions are recovered in simulations with a few thousand discrete loci (at least for the typical map lengths considered in the present analysis). This is evident in fig. 2 of Appendix S4 which shows how the average trait value in a finite population with *L* discrete loci approaches a limit as *L* becomes large.

More generally, re-examining classical adaptation or introgression scenarios in the polygenic setting is becoming increasingly important in light of recent evidence from genome-wide association studies (GWAS) that points towards the highly polygenic (and even ‘omnigenic’) nature of several traits (Boyle et al. 2017). These studies suggest a trait architecture in which a moderate number of strongly or moderately selected loci are embedded within a very large number of small-effect loci that are undetectable even in very high-powered GWAS but nevertheless explain a large fraction of the heritability of the trait. This is consistent with the common observation that quantitative trait loci (QTL) often ‘break up’ into multiple loci when investigated in more high-powered studies (Flint and Mackay 2009); fine mapping of individual genes also points towards multiple variants within each gene that contribute to trait variation (Stam and Laurie 1996). A related observation is that the fraction of heritability that can be attributed to individual chromosomes is proportional to the length of the chromosome (Yang et al. 2011); the heritability explained by various functional gene groups has also been found to be approximately proportional to the size of the group (Boyle et al. 2017). All of these suggest a very large number of trait loci spread more or less uniformly across the genome.

A second observation concerns the large number of deleterious recessive mutations present in the genomes of most species; these mutations are characterised by a wide range of selective effects, but with an average effect that is quite small (approximately 0.1% for Drosophila, see Charlesworth (2015)). Both these lines of evidence point to the relevance of the infinitesimal model with linkage as a way of understanding polygenic evolution. This model is of course an abstraction of reality, but it is arguably no less realistic than the usual assumption that the effects identified by GWAS or by QTL mapping are concentrated at single loci. A key question for the field is whether the effects on complex traits are typically due to single variants, or rather, are typically the combined effect of very many linked loci. Ideally, data would be tested against the alternative extremes, of a single locus vs. effects distributed over some extended region.

## SUPPORTING INFORMATION FOR: Introgression of a block of genome under infinitesimal selection

## Appendix S1 Evolution of trait values in the source population under the infinitesimal model with linkage

We describe below how the distribution of trait values in an infinite population evolves under the infinitesimal model with linkage. For simplicity, we consider a panmictic population and assume that there is no selection on the trait. Then, the frequency *P*(*z*_1_, *z*_2_) of diploid individuals with haplotypes associated with trait values *z*_1_ and *z*_2_, is simply the product of the haploid or gametic frequencies, i.e., *P*(*z*_1_, *z*_2_)=*P*(*z*_1_)*P*(*z*_2_).

The haplotype frequencies change from one generation to the next according to:

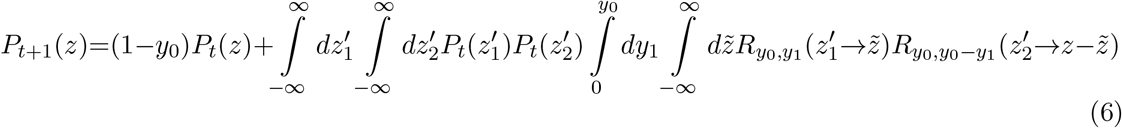

The first term on the right hand side represents the probability that there is no recombination within the genomic region contributing to trait value, so that the gamete has the same trait value as one of the haplotypes present in the diploid parent. The second term describes generation of a gamete by recombination between parental haplotypes with trait values 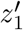 and 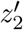. The map lengths of blocks inherited by the gamete from the two haplotypes are *y*_1_ and *y*_0_‒*y*_1_ (where *y*_1_ is uniformly distributed in [0, *y*_0_]); the additive contributions of these blocks is 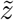 and 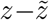 respectively. The term 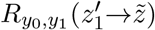 represents the probability that given a block of map length *y*_0_ and trait value *z*_1_, a daughter block of length *y*_1_ has an associated contribution 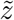 (see eq. 1 in the main text).

For a population in equilibrium, we expect *P*_*t*+1_(*z*)=*P*_*t*_(*z*). Approximating the equilibrium *P*(*z*) by a normal distribution with mean 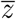 and variance *V*, and substituting into eq. (6), yields *V*=*V*_0_*y*_0_. The equilibrium variance of trait values in diploid individuals is thus 2*V*_0_*y*_0_ in the absence of selection.

If there is selection on the trait, then it is necessary to consider the evolution of diploid frequencies, since selection tends to build up correlations between the trait values associated with the two haplotypes carried by an individual. Moreover, eq. 1 in the main text does not describe inheritance of trait values in situations where there is LD between tightly linked regions of the genome (for instance, due to stabilizing selection, see (Lande 1976)). Extending the infinitesimal model to a scenario with selection and linkage is an interesting direction for future work.

## Appendix S2 Moments of the number of descendants and total amount of introgressed genome under the branching process approximation

For simplicity, we consider below the distribution of the number *N_t_*|_*y*,*z*_ of descendants of an introduced block with initial map length *y* and trait value *z*. Recursions for the Laplace transform of this distribution can be obtained similarly to eq. 3 of the main text, and are given by:

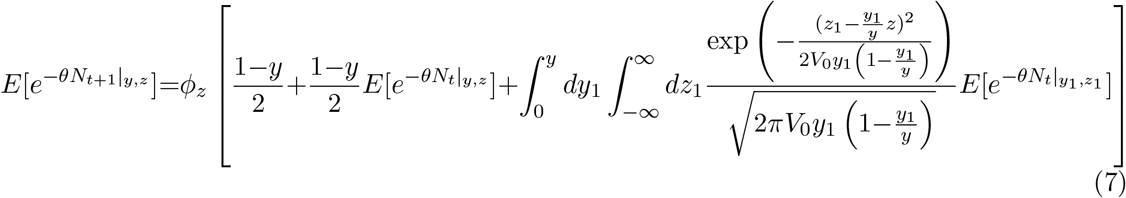

Differentiating the above equation with respect to *θ* and using *ϕ_z_*[*f(θ)*]=exp[−2*e^βz^*(1−*f(θ)*] (where *f(θ)* is the term in the square brackets above) yields the following recursions for the first two moments of the block number distribution.

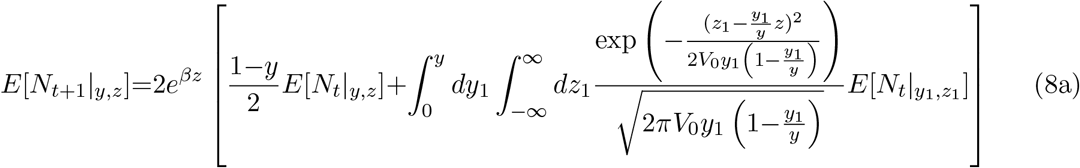

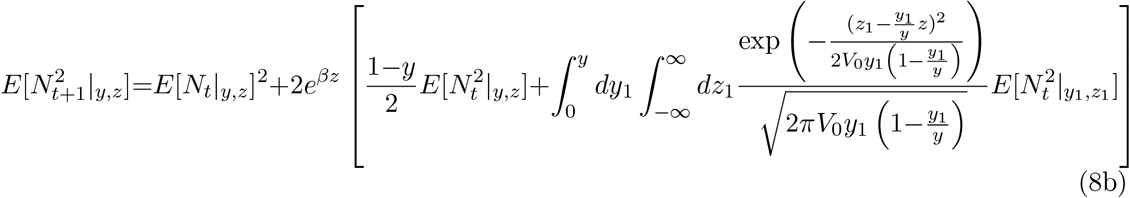

with the initial condition 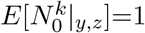 for all *k*. Note that eq. (8a), which gives recursions for the average number of descendants, is actually exact since the first moment of the distribution of *N* is insensitive to the correlations in trait values between different individuals. Higher moments, however, are affected by these correlations; thus eq. (8b) which gives recursions for 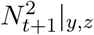 or alternatively for the variance in the number of descendants can only be written under the branching process approximation.

The distribution of the total amount of introgressed genetic material *L_tot_*(*t*) surviving at time *t*, obtained by summing over the map lengths of all descendant blocks of the introduced haplotype, can be obtained in a similar way. The distribution and the moments follow identical recursions as *N*, but the initial condition is now 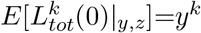 for all *k*>0. As for eq. 3 in the main text, we can numerically iterate eqs. (8a) and (8b) by we discretize (*y,z*) space and replace the two-dimensional integral by a double summation.

In principle, it is also possible to compute the *average* (and not just the total) length of surviving blocks from the joint distribution of *L_tot_* and *N* at any time *t*, which can be obtained by inverting the joint moment generating function *E*[*e*^−*θN*(*t*)−*λL_tot_*(*t*)^]_*y*,*z*_, but this is computationally quite laborious.

## Appendix S3 Effective parameters governing introgression under the infinitesimal model with linkage

To describe the continuous time dynamics for the number of descendants of an introduced block, we re-express eq. (8a) in terms of *β→βδt* and *y*→*yδt*, and further write *y* as *y*=*rL* where *r* is the recombination rate and *L* is the physical length of the block of interest. We also use the genic variance per unit physical length *σ*^2^, which is related to the genic variance per unit map length as *V*_0_*y*=*σ*^2^*L*. This yields:

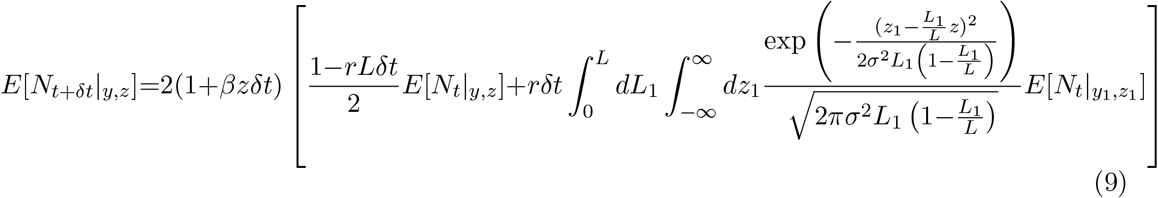

Taking the limit *δt*→0 and retaining the lowest order term in *δt* gives:

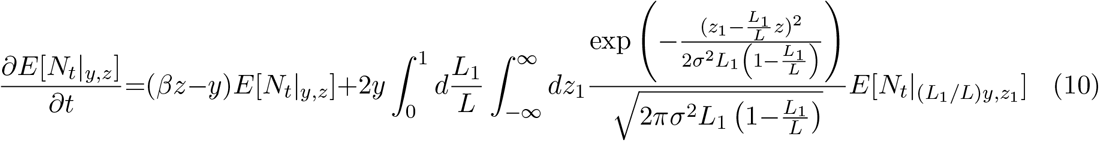

If we now denote *L*_1_/*L* by *α*, and divide the above equation throughout by *y*, we obtain eq. 4 in the main text, with 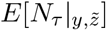 re-written as 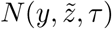. Equation 4 is expressed in terms of map length *y*, the scaled trait value 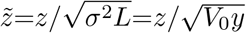, the ratio of selection to recombination strength 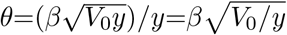, and rescaled time *τ*=*yt*.

Recall that at τ=0, we have 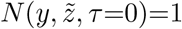. Thus the number of descendants after an infinitesimal interval *δτ* must be 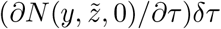 which is a function of θ, 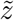 and *δτ*, but independent of *y*. If we now make the ansatz 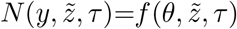 then it follows from eq. 4 in the main text that 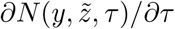 is also independent of *y*, so that 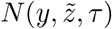 is independent of *y* at all later times. A similar argument can be made for the expected value *L_tot_* of the total amount of introgressed genome, which satisfies the same equation as eq. 4 (in the main text), but with the initial condition 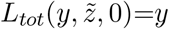; this then yields the solution 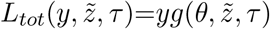. Note that 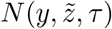 and 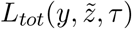 here refer to the unconditional expectation values, i.e., they have not been normalized by the probability that at least some part of the introduced block survives.

We test the validity of these scaling arguments for the true model of introgression into an infinite native population (which makes no assumptions about correlations between trait values of descendants). This model involves discrete generations and does not neglect correlations between trait values of different descendants of a block (see main text). Figure 9(a) shows the average number of descendants 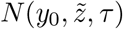 of a block with map length *y*_0_ and rescaled trait value 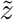 as a function of rescaled time *τ*=*yt*, for various values of *θ* and 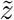 in the true model. For each 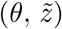 combination, we simulate introgression for three values of map length: *y*_0_=0.04 (red), *y*_0_=0.16 (green) and *y*_0_=0.36 (black), and find that there is perfect scaling collapse of 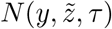 versus rescaled time τ for blocks of different lengths, as long as *z* and *β* are varied accordingly to hold 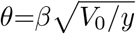 and 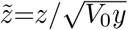 constant. We also find excellent scaling collapse for 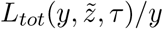 versus *τ* for different values of *y* (fig. 9(b)), in accordance with the scaling arguments above.

**Figure 9:**
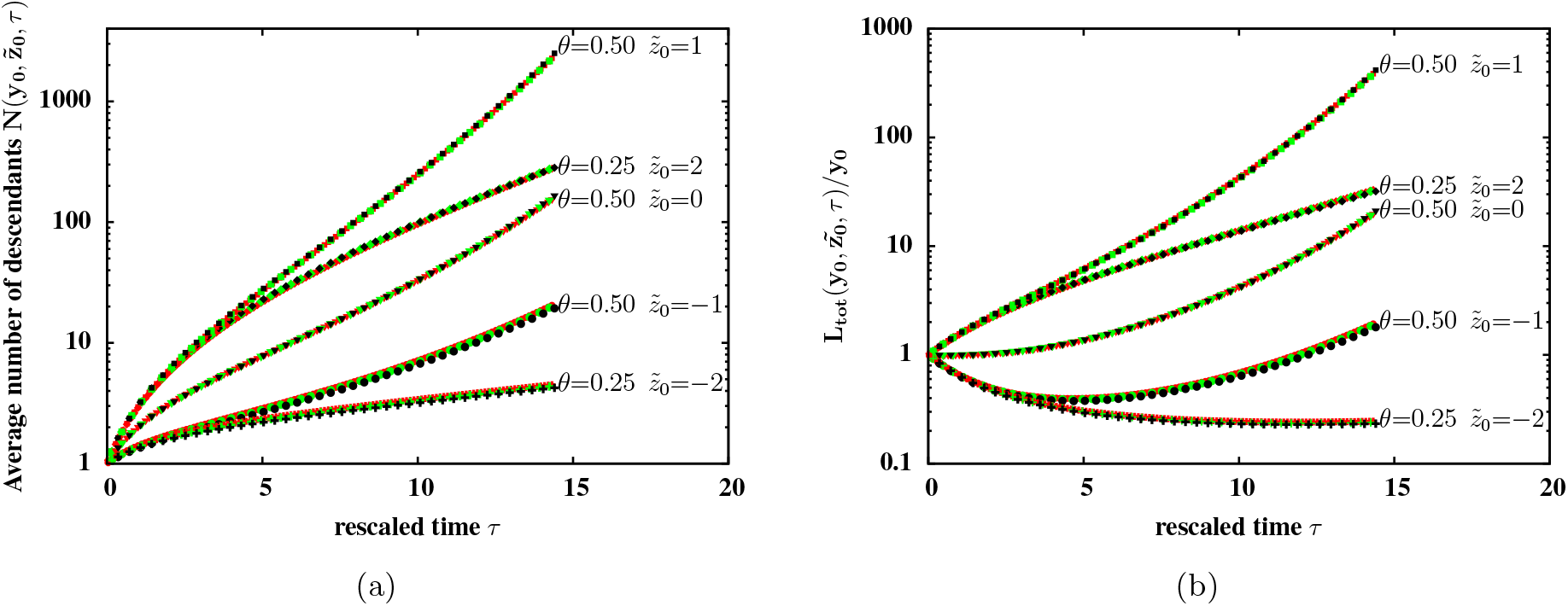
(a) Average number 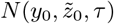 of descendants of an introduced block of map length *y*_0_ and effective trait value 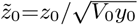 at rescaled time *τ*, vs. *τ* for three values of map length *y*_0_=0.04 (red), *y*_0_=0.16 (green) and *y*_0_=0.36 (black) and various 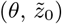 combinations (specified on the plots). (b) Average total amount 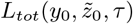 of introgressed genome (in units of *y*_0_, the map length of the selected region) vs. *τ*. For each 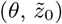 combination, the 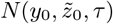 vs. *τ* curves (or the 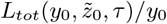 vs. *τ* curves) corresponding to the three values of *y*_0_ coincide completely.

Note that the *θ*=0.5, 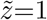 curve and the *θ*=0.25, 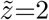 curve do not coincide although they are both characterised by the same value of 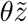. This illustrates an important distinction between the infinitesimal framework and an alternative model (also obtained by taking the *V*_0_→0 limit of the infinitesimal model) in which the trait value of the offspring is deterministic and simply proportional to the length of the block inherited from the parent (Barton 1983). In the latter model, it is the composite parameter 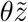 that governs introgression dynamics.

## Appendix S4 Individual-based simulations with discrete loci

Figure 4 of the main text shows results of individual-based simulations of finite populations with genome blocks containing *L*=2^12^ uniformly-spaced, discrete loci. We carry out simulations with various values of *L* and find that most quantities of interest approach a limit as *L* becomes large. This is demonstrated in fig. 10 for the average trait value in a population of size *N*=1000. Figure 10 also shows that the long-term advance under selection is higher when the same trait value (here *z*_0_=1) is due to a larger number of loci with a smaller variance in additive contributions.

**Figure 10:**
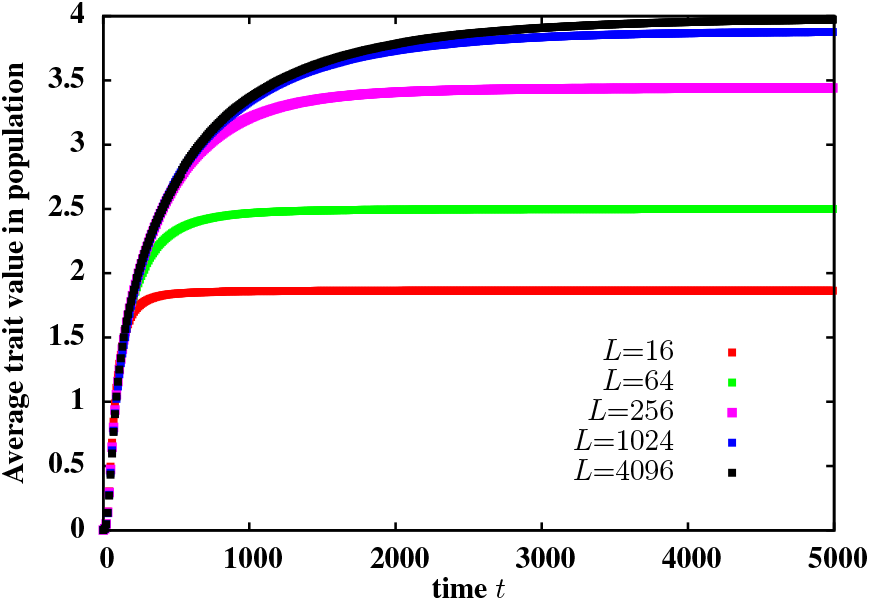
Average trait value associated with individuals in the population, versus *t*, for various values of *L*. Results are from individual-based simulations of a population of size *N*=1000, for a single introduced block with trait value *z*_0_=1 and map length *y*_0_=0.25, for selection strength 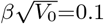.

## Appendix S5 Variability between replicates corresponding to the same Brownian path

Figure 1 in the main text illustrates the variability between two replicates involving introgression of two different genomes described by distinct Brownian paths (which are drawn from the same distribution, characterized by a particular value of *y*_0_, *z*_0_ and *V*_0_). These Brownian paths are likely to differ from each other by chance in various details, e.g., the location of the genomic fragment associated with the highest additive contribution, as well as the length and actual trait value associated with this fragment. Thus, some Brownian paths could be associated with a higher probability and rate of introgression, if they happen to contain a fragment that is particularly favourable.

However, there is additional variability arising from the random nature of recombination and reproduction events. This is illustrated in fig. 11 which shows results of replicate simulations for *the same* introduced block (corresponding to the same Brownian path). These replicates thus represent two different stochastic histories of birth and recombination events. Each column corresponds to one replicate, with the upper and lower panels depicting descendant individuals who carry some fragment of the introduced block, *t*=20 and *t*=80 generations after the initial hybridization event respectively. Note that there is significant variability between the two replicates (columns)— the sub-blocks that survive the initial, nearly-neutral phase of the introgression process are completely different in the two cases; they are also associated with different trait values, and hence proliferate at different rates. Consequently, the number of descendant blocks at *t*=80 is also quite different in the two cases.

**Figure 11:**
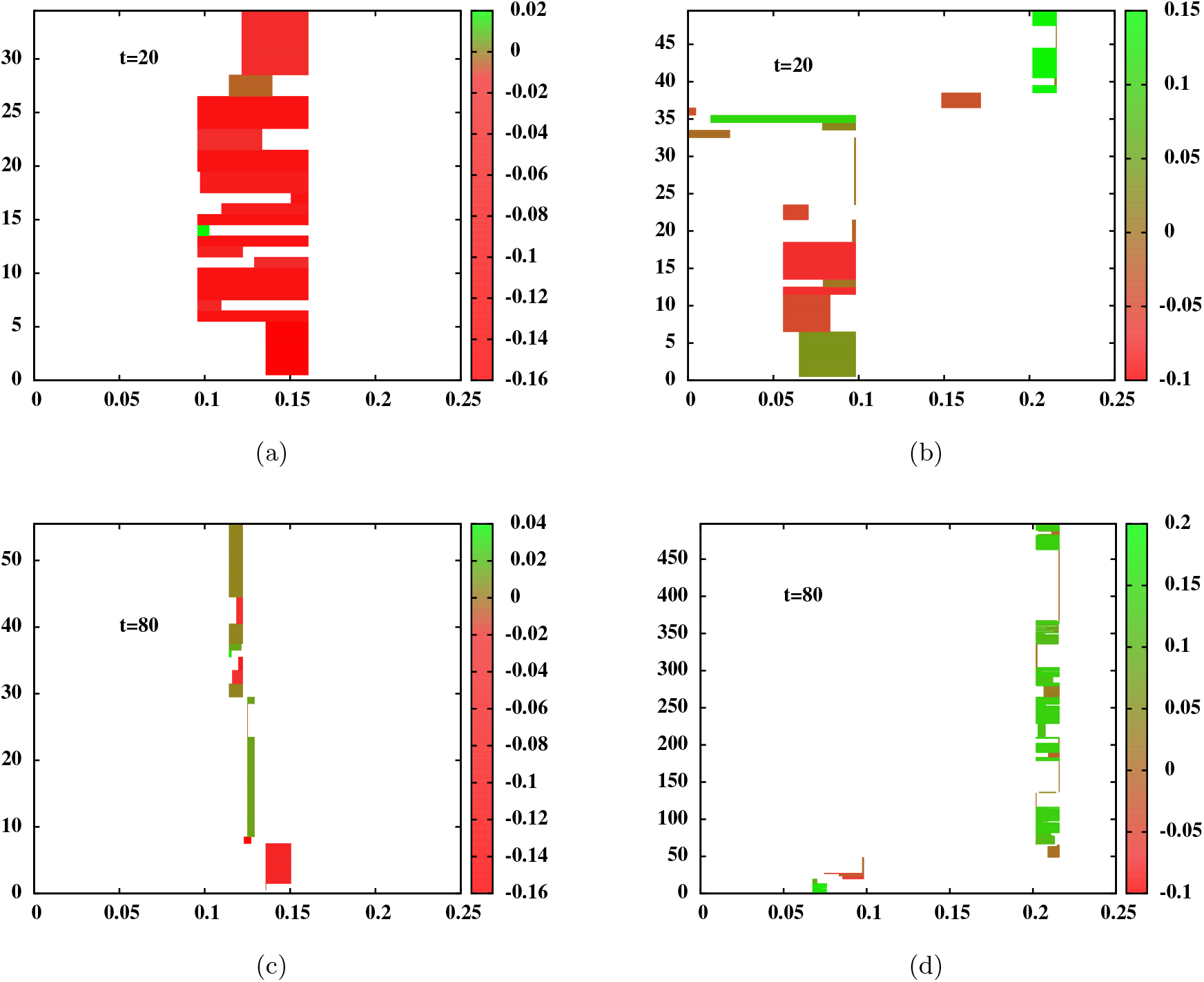
Snapshots of descendants of the introduced genome at *t*=20 (fig. 11(a)) and *t*=80 (fig. 11(c)) for a single realization of the introgression process for an initially neutral block (*z*_0_=0) with map length *y*_0_=0.25 and 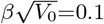. Figures 11(b) and 11(d) show the corresponding snapshots for a second (independent) realization for introgression of the same genomic block (represented by the same Brownian path as in the case of figs. 11(a) and fig. 11(c)). Each horizontal line represents the genome of an individual carrying a fragment of the introduced block; the coloured portion represents the introgressed fragment while the white portions represent native blocks. The trait value associated with each introgressed fragment is encoded by the colour of the block (green for positive trait values and red for negative values) and can be read off from the accompanying colour scale. The y-axis of each plot indexes the descendants carrying introgressed block fragments. The x-axis shows positions along the genomic region influencing trait value (here having map length *y*_0_=0.25).

This example thus suggests that for various parameter regimes, the stochasticity inherent in recombination and other evolutionary processes may be as (or even more) important a source of variability between replicates as the variability associated with different introduced blocks. We expect the former to be especially significant when segregating blocks are few, long, and associated with weakly non-zero effects (in which regime recombination is expected to influence dynamics more than selection). Understanding the consequences of different kinds of stochasticity during different phases of the introgression process is an interesting direction, and will be explored in more detail in a future study.

